# Xenophagocytosis blockade enhances interspecies chimerism

**DOI:** 10.1101/2025.10.14.682291

**Authors:** Sicong Wang, Kouta Niizuma, Daniel Dan Liu, Fabian P. Suchy, Hideyuki Sato, Ayaka Yanagida, Hideki Masaki, Masashi Miyauchi, Saman Tabatabaee, Nathan Hidajat, Joydeep Bhadury, Carsten T. Charlesworth, Jinyu Zhang, Irving L. Weissman, Hiromitsu Nakauchi

**Author notes:** Correspondence (H.N.). These authors contributed equally (Sicong Wang, Kouta Niizuma, Daniel Dan Liu).

## Abstract

Organ shortage remains a major challenge in transplantation medicine. Interspecies blastocyst complementation is a promising approach to generate human organs in livestock hosts. However, getting xenogeneic donor cells to engraft and expand at early stages remains challenging. Here we identify an innate immune barrier, wherein host macrophages selectively recognize and eliminate viable xenogeneic donor cells. These events represent a form of phagoptosis and highlight a xenogeneic clearance process that we term xenophagocytosis. We identify the mechanism by which host macrophages selectively phagocytize xenogeneic donor cells: xenogeneic cells display elevated phosphatidylserine, an “eat-me” signal recognized by host macrophages through phagocytic receptor Axl. Xenophagocytosis blockade improves both rat and human donor chimerism in mouse embryos, indicating a conserved mechanism. These findings reveal potential mechanisms by which innate immune cells eliminate xenogeneic cells in early embryogenesis to preserve species integrity and offer improved strategies for generating human organs in livestock.

## INTRODUCTION

Finding solutions to the critical shortage of donor organs is imperative^1^. Patients suffer from lifelong detrimental effects of immunosuppression after receiving allogeneic or xenogeneic organ transplants^2^, which highlights the importance of generating autologous organs. Inspired by the fact that successful engraftment of hematopoietic stem cells requires the creation of empty niches through myeloablation^3,4^, we developed the concept of an “organ niche” and established a method to generate pluripotent stem cell (PSC)-derived organs within interspecies chimeric animals^5–9^.

Under this concept, we successfully generated a rat pancreas within a mouse host^5^. Furthermore, in a reciprocal experiment, we generated a mouse PSC-derived pancreas within a rat host and demonstrated that islets isolated from these complemented pancreata could cure diabetes in mice without the need for immunosuppression^7^. Through these studies, we observed that interspecies chimeras exhibited significantly lower birth rates and donor cell chimerism compared to intraspecies chimeras^5,10^. Additionally, time-course analysis of rat > mouse (rat donor cells injected into mouse host embryos) interspecies chimeric embryos revealed a marked decrease in donor-derived cells between embryonic days E9 and E11^10^. We attribute the restricted engraftment and development of xenogeneic cells arising at this stage to a “xenogeneic barrier.”

The nature of the xenogeneic barrier remains elusive and multifactorial. Although developmental incompatibilities—such as mismatches in donor-host developmental stages^11–14^, adhesion molecules^15–17^, developmental tempo^18^, cell fitness levels^19^, or signaling pathways^20,21^—have been proposed and investigated^22–25^, the immunological incompatibility during embryogenesis remains largely unexplored.

Given that the emergence of the earliest immune cells^26,27^ roughly coincides with the timing of the xenogeneic barrier, as well as the critical role of immune interactions in xeno-transplantation biology^28,29^, we hypothesize that immunological incompatibility represents one major, yet overlooked, component of the xenogeneic barrier.

Here we investigated the role of immune cells in the development of interspecies chimeric embryos between mouse and rat. We discovered that the earliest innate immune cell type, primitive macrophages^26^, preferentially recognize xenogeneic donor cells and eliminate them through phagocytosis. We show that the mechanism of macrophage recognition is independent of apoptotic cell clearance^30^; instead, it is due to the imbalance between “eat-me” and “don’t eat me” signals displayed on the xenogeneic donor cells. Through genetic depletion of macrophages from host embryos, or overexpression of species matched “don’t eat-me” signal on the donor PSC, we achieve increased chimerism in multiple organs.

## RESULTS

### Xenophagocytosis in interspecies chimeric embryos

To define the developmental stage at which xenogeneic donor cells begin to be actively eliminated, we generated rat > mouse chimeras by injecting EGFP labeled rat pluripotent stem cells into mouse blastocysts followed by embryo transfer. Donor chimerism in the whole embryo was analyzed by digital PCR at E9.5, E11.5 and E14.5^31^. As described previously^10^, although high chimerism embryos could be observed at E9.5 (mean chimerism: 21.9%), only low chimerism embryos were recovered at E11.5 afterwards (mean chimerism: 2.8%) (Figure 1A). This sharp decline in chimerism suggests that donor rat cells were actively eliminated by host mouse cells starting around E9.5. Notably, this stage is prior to the establishment of the acquired immune system by T and B cells^32^, yet coincides with the period when primitive macrophages emerge^26,27^.

**Figure 1.**
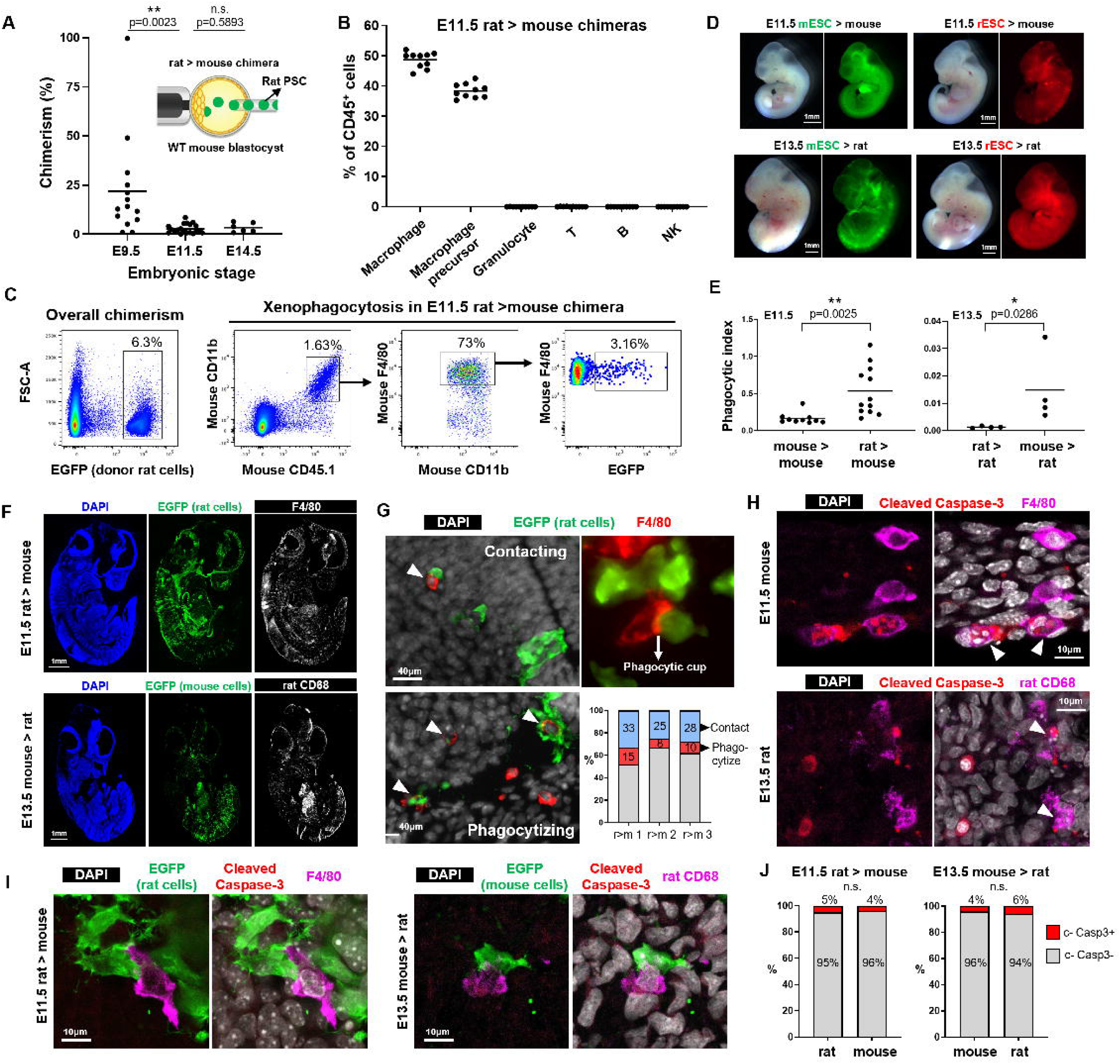
Recipient embryonic macrophages eliminate xenogeneic donor cells through an apoptosis-independent mechanism. **(A)** Whole embryo chimerism of rat > mouse chimeras at three embryonic stages. Each dot represents one chimeric embryo. The black bar indicates the mean chimerism of each stage. Chimerism was quantified by ddPCR. n = 14 (E9.5), 20 (E11.5), 6 (E14.5). Statistical significance was assessed using unpaired two tailed t-test. n.s., not statistically significant; *p < 0.05, **p < 0.01. **(B)** Immune cell profiling of E11.5 rat > mouse chimeras. Mean percentage of each immune cell type over mouse CD45^+^ cells is shown. Each dot represents one chimeric embryo. n = 10 chimeras. NK, natural killer cells. **(C)** Quantification strategy to identify “eating macrophages” in E11.5 rat > mouse chimeras using flow cytometry. Mouse macrophages were defined as mCD45.1^+^ mCD11b^+^ F4/80^+^ cells. Among these, EGFP^+^ macrophages are classified as “eating macrophages”. A distinct population of homogeneous, EGFP^^high^ cells lacking mouse macrophage markers was defined as intact donor rat cells and used for quantifying chimerism. **(D)** Representative images of intraspecies (mouse > mouse, rat > rat) and interspecies (rat > mouse, mouse > rat) chimeras at morphology matched stages. E11.5 mouse embryos morphologically match with E13.5 rat embryos. Both bright field and fluorescent pictures are shown. Donor mouse ESCs are labelled with EGFP and donor rat ESCs are labelled with tdTomato. **(E)** Comparison of the phagocytic index across intraspecies (mouse□>□mouse, rat□>□rat) and interspecies (rat□>□mouse, mouse□>□rat) chimeras. The phagocytic index was calculated as the percentage of mCD11b□F4/80□ macrophages containing EGFP□ donor signal, normalized to overall donor chimerism. Data are shown for E11.5 chimeras (n□=□10 for mouse > mouse, n = 12 for rat > mouse) and E13.5 chimeras (n□=□4 for rat > rat and n = 4 for mouse > rat). Each dot represents one chimeric embryo. Graph shows mean phagocytic index. Statistical significance was assessed using unpaired two tailed t-test (left panel) or unpaired Mann-Whitney U non-parametric test (right panel). n.s., not statistically significant; *p < 0.05, **p < 0.01. **(F)** Representative fluorescent images of sagittal sections from E11.5 rat□>□mouse and E13.5 mouse□>□rat chimeras. Sections were stained for F4/80 (white, host mouse macrophages), CD68 (white, host rat macrophages), DAPI (blue) and EGFP (green, donor cells). Macrophages are widely distributed throughout the embryonic tissues at both stages. **(G)** Representative fluorescent images of F4/80□ mouse macrophages and EGFP□ rat donor cells in E11.5 rat□>□mouse chimeras. Two types of interactions are shown: “contacting” macrophages, which are in direct contact with donor cells, and “phagocytizing” macrophages, which contain internalized EGFP signal. Arrowheads indicate macrophages in each interaction state. A phagocytic cup structure is visible in the right panel. Percentage of “contacting” and “phagocytizing” macrophages from three E11.5 rat > mouse chimeras (r>m 1-3) was quantified. **(H)** Representative fluorescent images of E11.5 mouse and E13.5 rat embryos showing macrophages clearing apoptotic cells. Cleaved caspase-3□ signals (red) indicate apoptotic cells, often co-localized with fragmented nuclear DNA visualized by DAPI (white). These apoptotic remnants are located within F4/80□ (mouse) or CD68□ (rat) macrophages. White arrowheads mark representative macrophages containing engulfed apoptotic material. **(I)** Representative fluorescent images showing xenogeneic phagocytosis in interspecies chimeras. Left, an F4/80□ mouse macrophage in an E11.5 rat□>□mouse chimera engulfs an EGFP□ rat cell lacking cleaved caspase-3 signal. Right, a CD68□ rat macrophage in an E13.5 mouse□>□rat chimera engulfs a non-apoptotic EGFP□ mouse cell. **(J)** Quantification of apoptosis status for cells in contact with host macrophages in E11.5 rat > mouse chimeras (left) and E13.5 mouse > rat chimeras (right). Rat and mouse cells in interspecies chimeras are plotted separately. Statistical significance was assessed using Fisher’s exact test. n.s., not statistically significant; *p < 0.05. n = 214 (rat, left), n = 240 (mouse, left), n = 126 (mouse, right), n = 155 (rat, right). c-Casp3: cleaved caspase-3. See also Figure S1.

To characterize the host immune cell populations (CD45^+^) present at this critical stage in chimeras, we performed comprehensive immune profiling of E11.5 rat > mouse chimeras by flow cytometry. We detected macrophages (CD11b^+^F4/80^+^) and macrophage precursors (CD11b^+^F4/80^−^), but not other immune cell types (Figures 1B and S1A). This immune profile aligns well with previous studies demonstrating that primitive macrophages are the predominant immune cell type present at this developmental stage^33^, preceding the emergence of adaptive immune cells^32^. Based on these observations, and considering the previously proposed concept of phagoptosis^34,35^—a form of cell death in which viable cells are eliminated by phagocytes—we hypothesized that primitive macrophages might selectively recognize and remove xenogeneic donor cells, thereby contributing to the observed decline in chimerism.

To directly test whether host macrophages actively phagocytize the donor rat cells, we focused on the macrophage population (CD11b^+^F4/80^+^) from E11.5 rat > mouse chimeras. Within this population, we identified a clear subset of mouse macrophages that were EGFP positive (Figure 1C). Imaging flow cytometry analysis^36^ of this subset revealed that the EGFP signal was localized as discrete intracellular puncta within macrophages, confirming genuine engulfment of EGFP-labeled rat cells rather than cell doublets or aggregates (Figure S1B). We term this phenomenon “xenogeneic phagocytosis,” or “xenophagocytosis” for short.

To examine whether xenophagocytosis also occurs in other host species, we generated mouse > rat chimeras by injecting EGFP-labeled mouse PSCs into rat blastocysts. Since rat embryos develop two days slower than mouse embryos around mid-gestation stages, we focused on comparing E13.5 rat embryos with E11.5 mouse embryos in this study. Flow cytometry analysis of E13.5 mouse > rat chimeras similarly revealed rat CD11b^+^ macrophages containing EGFP signals, indicating that xenophagocytosis is not unique to mouse hosts (Figure S1C). Since phagocytosis is a common physiological process during embryonic development, we next asked whether host macrophages preferentially target xenogeneic donor cells over species-matched donor cells. To address this, we systematically generated stage-matched intra- and interspecies chimeras of all four combinations between mouse and rat donors and hosts and quantitatively compared macrophage phagocytic activity by flow cytometry (Figures 1D and S1C). We defined a “phagocytic index” as the percentage of host macrophages containing donor-derived EGFP signals normalized to the chimerism of each chimera, as this accounts for the variability in chimerism from embryo to embryo. Notably, interspecies chimeras (rat > mouse and mouse > rat) exhibited significantly higher phagocytic indices compared to intraspecies chimeras (mouse > mouse and rat > rat), which demonstrated that macrophages preferentially eliminate xenogeneic donor cells over species-matched cells (Figure 1E).

To spatially characterize the interactions between xenogeneic donor cells and host macrophages, we performed immunostaining for macrophage markers (F4/80 for mouse macrophages in E11.5 rat > mouse chimeras, and rat CD68 for rat macrophages in E13.5 mouse > rat chimeras). At these developmental stages, macrophages were observed throughout the embryonic tissues (Figure 1F). In E11.5 rat > mouse chimeras, we identified two distinct types of interactions between mouse macrophages and EGFP-labeled rat donor cells: direct cell-cell contact without internalization (“contacting”) and active engulfment of donor cells (“phagocytizing”). 28.7 ± 4.1% of macrophages were found in direct contact with rat donor cells without internalization (Figures 1G and S1D). In addition, “contacting” macrophages with phagocytic cup structures, indicative of the initial stage of phagocytosis, were frequently observed (Figure 1G). 11 ± 3.6% of macrophages contained EGFP signal clearly localizing within the cytoplasm, representing cells in the digestion phase of phagocytosis.

To determine whether macrophages actively eliminate viable xenogeneic donor cells or merely clear cells already undergoing apoptosis, we assessed apoptosis status by immunostaining for cleaved caspase-3^37^. Distinguishing between these two possibilities is critical, as active elimination of viable donor cells would indicate macrophages as direct mediators of the xenogeneic barrier, rather than passive scavengers of dying cells. Most cleaved caspase-3-positive signals co-localized with fragmented DNA and were found interior to host macrophages, consistent with the canonical role of macrophage-mediated clearance of apoptotic cells (Figure 1H) ^38–40^. However, the majority of xenogeneic donor cells interacting with host macrophages were negative for cleaved caspase-3 and lacked fragmented DNA morphology (Figures 1I and 1J). This apoptosis-negative state of xenogeneic donor cells was consistently observed in both rat > mouse and mouse > rat chimeras, suggesting that macrophages predominantly target viable xenogeneic donor cells, instead of merely scavenging already-dying donor cells (Figures 1I and 1J). This pattern of phagocytosis closely resembles a form of primary cell death known as phagoptosis, in which viable cells are eliminated by phagocytes without undergoing apoptosis^34,35^. Our findings suggest that xenophagocytosis may represent a physiological instance of phagoptosis occurring in the embryonic development of interspecies chimeras.

### Macrophage deficient host embryos exhibit increased xenogeneic donor chimerism

Based on these observations, we hypothesized that host macrophages actively recognize and eliminate xenogeneic donor cells through xenophagocytosis during mid-gestation, resulting in the sharp decline of chimerism observed from E9.5 onward (Figure 1A). To directly test this hypothesis, we disrupted macrophage development in host embryos by knocking out PU.1 (gene name: Spi1), a master transcription factor essential for myeloid lineage specification during primitive hematopoiesis^41–43^.

We electroporated ribonucleoprotein (RNP) complexes consisting of Cas9 protein and sgRNA targeting mouse Spi1 into fertilized mouse zygotes (Figure 2A)^44^. EGFP-labeled rat PSCs were then injected into genome-edited or control mouse blastocysts. Whole-embryo donor chimerism was quantified at three developmental stages (E9.5, E11.5, and E14.5) using droplet digital PCR (ddPCR) assay^31,45^. PU.1 knockout host embryos showed significantly increased rat donor chimerism at E11.5 and E14.5 compared to control embryos (Figure 2B). In contrast, no significant difference of donor chimerism was observed at E9.5, consistent with the timing of macrophage colonization and expansion, which begins around E9.5 and is still limited at this early stage^27^.

**Figure 2.**
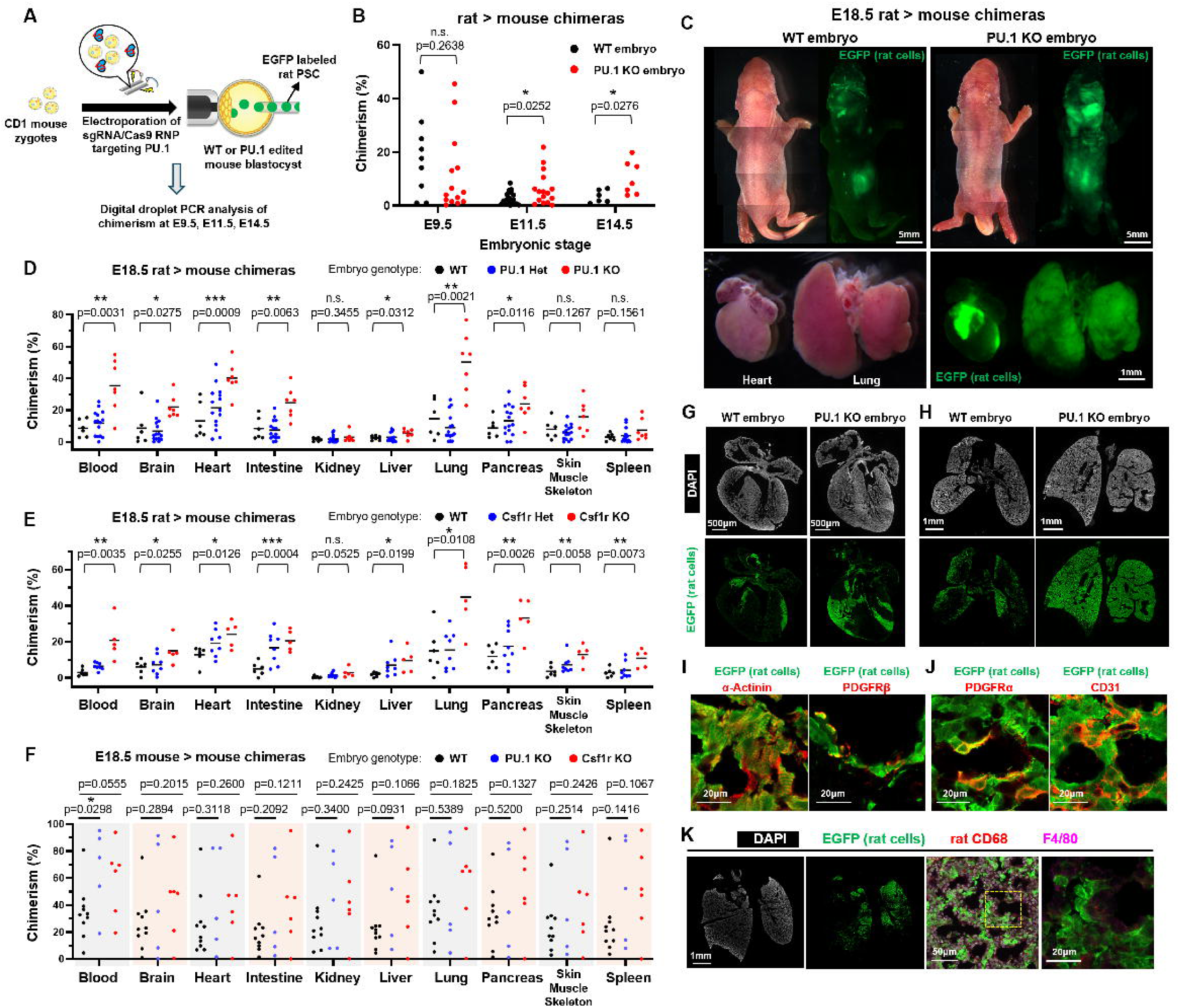
Genetic ablation of host embryo macrophage increases xenogeneic donor chimerism across multiple organs. **(A)** Schematic of rat□>□mouse chimera generation using mouse embryos with PU.1 gene disruption. For E9.5, E11.5, and E14.5 analysis, PU.1 was knocked out by electroporating Cas9/gRNA RNP into wild-type mouse zygotes prior to blastocyst injection. For E18.5 analysis, PU.1-deficient embryos were obtained by intercrossing PU.1 heterozygous mice. **(B)** Whole embryo chimerism analysis of rat > mouse chimeras at three different embryonic stages. The genotype of mouse embryos is either wild type or PU.1 knockout. Each dot represents one chimeric embryo. Chimerism was quantified by ddPCR. For WT embryos, n = 9 (E9.5), 20 (E11.5), 6 (E14.5). For PU.1 knockout embryos, n = 14 (E9.5), 16 (E11.5), 7 (E14.5). Statistical significance was assessed using unpaired two tailed t-test. n.s., not statistically significant; *p < 0.05. **(C)** Representative bright-field and fluorescence images of E18.5 rat□>□mouse chimeras using WT or PU.1 knockout mouse embryos. EGFP indicates xenogeneic donor rat cells. Bottom panels show dissected heart and lung from a rat > PU.1 knockout mouse chimera with high donor chimerism. **(D)** Donor chimerism in ten organs from E18.5 rat□>□mouse chimeras with WT, PU.1 heterozygous, or PU.1 knockout mouse hosts. Each dot represents one organ dissected from one chimera. Black bars indicate group means. Chimerism was quantified by ddPCR. n = 6 (WT), 15 (heterozygous), and 7 (KO). Statistical significance was assessed using unpaired two tailed t-test. n.s., not statistically significant; *p < 0.05, **p < 0.01, ***p < 0.001. **(E)** Donor chimerism in ten organs from E18.5 rat□>□mouse chimeras with WT, Csf1r heterozygous, or Csf1r knockout mouse hosts. Each dot represents one organ dissected from one chimera. Black bars indicate group means. Chimerism was quantified by ddPCR. n = 6 (WT), 8 (heterozygous), and 5 (KO). Statistical significance was assessed using unpaired two tailed t-test. n.s., not statistically significant; *p < 0.05, **p < 0.01, ***p < 0.001. **(F)** Donor chimerism in ten organs from E18.5 mouse□>□mouse chimeras with WT, PU.1 knockout, or Csf1r knockout mouse hosts. Each dot represents one organ dissected from one chimera. Black bars indicate group means. Chimerism was quantified by ddPCR. n = 10 (WT), 5 (PU.1 KO), and 6 (Csf1r KO). Statistical significance was assessed using unpaired two tailed t-test. n.s., not statistically significant; *p < 0.05. **(G-H)** Representative heart (G) and lung (H) sections from E18.5 rat□>□mouse chimeras stained for EGFP□ donor rat cells. Host mouse embryos were either WT or PU.1 knockout. **(I)** Magnified view of heart tissue sections from an E18.5 rat > PU.1 KO mouse chimera stained for EGFP along with lineage-specific markers: α-Actinin (cardiomyocytes), PDGFRβ (pericytes). **(J)** Magnified view of lung tissue sections from an E18.5 rat > PU.1 KO mouse chimera stained for EGFP along with lineage-specific markers: PDGFRα (mesenchyme), CD31 (endothelial cells). **(K)** Representative lung tissue section from an E18.5 rat > PU.1 KO mouse chimera stained for EGFP□ donor rat cells and species-specific macrophage markers: F4/80 (mouse) and rat CD68 (rat). See also Figure S2.

To obtain genetically defined internal controls and avoid potential confounding off-target effects from CRISPR genome editing, we next generated chimeras using mouse embryos collected from PU.1 heterozygous intercrosses. Homozygous knockout of PU.1 resulted in nearly complete loss of CD45^+^ immune cells at E11.5 and E18.5, and the E18.5 embryos exhibited a rudimentary thymus, consistent with previously reported phenotypes^41,42^ (Figures S2A-S2D). We then generated chimeras as before, by injecting EGFP-labeled rat PSCs into mouse blastocysts obtained from PU.1^+/−1bp^ intercrosses, allowing subsequent genotyping to classify host embryos as wild-type, heterozygous, or homozygous knockout. We analyzed donor chimerism of ten organs in E18.5 chimeras. Chimeras with high donor chimerism in skin, heart, intestine, and lung were readily identified by strong EGFP fluorescence (Figure 2C), and genotyping revealed these embryos as PU.1 homozygous knockouts. Systematic analysis of organ chimerism from 28 chimeras revealed significantly increased donor chimerism in multiple organs of PU.1 homozygous knockout embryos compared to wild-type controls (Figure 2D). The greatest increases in chimerism were observed in blood (from 9% to 33%), heart (14% to 38%), intestine (8% to 26%), and lung (18% to 45%) (Figure 2D). Remarkably, we achieved donor chimerism levels as high as 55% in blood, 36% in brain, 57% in heart, 41% in intestine, and 77% in lung, substantially surpassing previously reported levels in rat > mouse interspecies chimeras^10,45^. No significant difference in chimerism was observed between PU.1 heterozygous and wild-type embryos, highlighting that complete loss of PU.1 function is required to enhance xenogeneic donor chimerism.

PU.1 is essential for the development of multiple immune lineages, including both lymphoid and myeloid cells, during primitive and definitive hematopoiesis^41–43^. To further clarify whether the increased xenogeneic chimerism observed in PU.1 knockout embryos was primarily due to macrophage depletion, we next targeted Csf1r, a gene specifically required for macrophage development and survival^38,46^. Homozygous Csf1r knockout neonates can be readily identified by their toothless phenotype and reduced macrophages in several organs (Figures S2E-S2G). Like before, we crossed Csf1r^+/−^ mice to obtain blastocysts with three potential genotypes (wild-type, heterozygous, or homozygous knockout), into which we injected rat PSCs. Organ chimerism analysis again revealed significantly increased rat donor chimerism in blood, heart, intestine, and lung of Csf1r knockout embryos compared to wild-type or heterozygous littermates (Figure 2E). The consistent increase in xenogeneic chimerism observed in both PU.1 and Csf1r knockout embryos strongly supports the conclusion that macrophage depletion in host embryos is a primary factor that enhances xenogeneic donor chimerism.

Our earlier observations indicated that xenophagocytosis differs from the apoptotic cell clearance occurring during embryonic development. Therefore, we hypothesized that macrophage depletion might specifically enhance interspecies chimerism, but not intraspecies chimerism. To directly test this hypothesis, we generated mouse > mouse chimeras by injecting EGFP-labeled mouse PSCs into PU.1 or Csf1r knockout mouse blastocysts. Organ chimerism analysis at E18.5 revealed no significant increase in donor chimerism in either PU.1 or Csf1r knockout embryos compared to wild-type controls (Figure 2F). Chimerism levels were highly variable among individuals in all three groups, consistent with the known variability of intraspecies mouse > mouse chimeras^31^. Together, these results demonstrate that genetic ablation of host macrophages significantly enhances xenogeneic donor (rat > mouse) chimerism in multiple organs but does not affect intraspecies (mouse > mouse) chimerism. This finding highlights xenophagocytosis as a unique immunological mechanism restricting interspecies chimerism.

We further analyzed chimeric heart and lung tissues at cellular resolution by immunohistochemistry. Microscopy analysis of heart sections revealed broader distribution of EGFP^+^ rat donor cells in PU.1 knockout host embryos compared to wild-type embryos (Figure 2G). Importantly, lineage-specific markers demonstrated efficient differentiation of rat PSCs into α-Actinin^+^ cardiomyocytes and PDGFRβ^+^ pericytes within the chimeric heart (Figure 2I). These results highlight the potential of macrophage depletion to significantly enhance xenogeneic contribution to clinically relevant cell types, such as cardiomyocytes. Similarly, we observed widespread EGFP^+^ rat donor cells in PU.1 knockout embryos compared to wild-type controls in lung (Figure 2H). However, lineage analysis revealed a skewed contribution of rat cells predominantly toward CD31^+^ endothelial cells and PDGFRα^+^ mesenchymal cells (Figure 2J). In contrast, rat-derived E-cadherin^+^ /Nkx2.1^+^ lung epithelial cells were rarely detected. This lineage-specific difference in xenogeneic contribution suggests that the strength or nature of the xenogeneic barrier may vary significantly among different cell types within the same organ. Alternatively, it may suggest different levels of evolutionary divergence in the molecular and cellular processes required for the organogenesis of each organ.

Another alternative explanation for the increased xenogeneic chimerism observed in macrophage-deficient embryos is that the donor rat PSCs might differentiate into macrophages, thereby reconstituting the depleted host macrophage population. To test this possibility, we carefully examined macrophage populations in lung tissue of rat > mouse chimeras generated using PU.1 knockout mouse blastocysts. Immunostaining confirmed the complete absence of host-derived (F4/80^+^) mouse macrophages, validating the robust macrophage depletion phenotype of PU.1 knockout embryos (Figure S2H). Importantly, despite abundant rat donor cells present, we detected few rat-derived macrophages (EGFP^+^ rat CD68^+^) in the lung (Figures 2K and S2H). This indicates that donor rat PSCs did not efficiently differentiate into macrophages to replace the depleted host macrophages. Thus, the observed increase in xenogeneic chimerism is likely due to the absence of host macrophages, rather than macrophage reconstitution by donor cells.

### Host macrophages recognize xenogeneic donor cells through TAM receptors

Macrophages are innate immune cells that survey their environment by integrating ‘eat-me’ and ‘don’t eat me’ signals, allowing them to discriminate foreign cells from self^47,48^. We thus sought to investigate which molecular pathways are utilized by embryonic macrophages in their preferential phagocytosis of xenogeneic rat cells. Chimeric rat > mouse E9.5 and E11.5 embryos were dissociated and stained with a surface marker panel consisting of CD45.1, CD11b, F4/80, CD80, CD38, CD206, SIRPα, and Ly6c. mCD45.1^+^mCD11b^+^ mouse myeloid cells were index-sorted into 96-well plates for single cell RNA sequencing (scRNA-seq) using Smart-seq3^49^(Figures 3A and 3B). Three sub-populations were collected from this population: mF4/80^−^ (pre-macrophage), mF4/80^+^EGFP^−^ (non-eating macrophage), and mF4/80^+^EGFP^+^ (eating macrophage). For internal comparison, we also sorted peritoneal and bone marrow (BM) macrophages from 12-week adult mice. Index sorting records the mean fluorescence intensity (MFI) of each sorted cell, providing the exact surface marker profile (immunophenotype) of each single cell transcriptome^50^. The recorded EGFP signal also provides a ground-truth label for whether each macrophage has recently phagocytized a rat cell.

**Figure 3.**
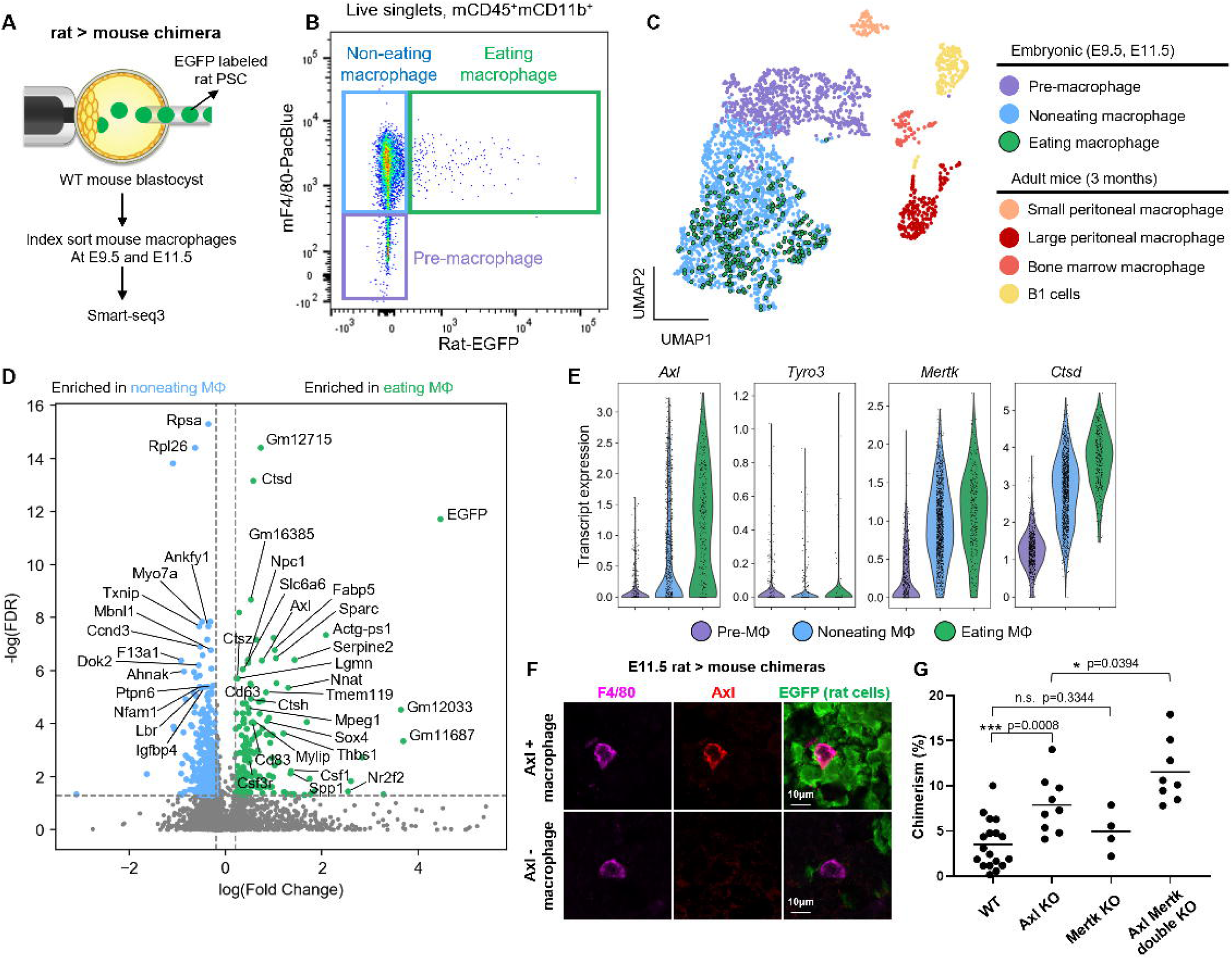
Single cell analysis of eating macrophages from interspecies chimeras identifies Axl as a receptor for xenogeneic cell recognition and phagocytosis. **(A)** Schematic of experimental setup. **(B)** Gating scheme for index sort. The macrophages from E9.5 and E11.5 rat > mouse chimeras are separated into three groups (Eating macrophages: F4/80^+^, EGFP^+^; Non-eating macrophages: F4/80^+^, EGFP^-^; Pre-macrophage: CD11b^+^, F4/80^-^). **(C)** UMAP of sorted myeloid cells. Macrophage subtypes are color coded on the UMAP. **(D)** Volcano plot of differential expressed genes between eating and noneating macrophages. Cutoff for log_10_ (FDR): 0.05, for log_10_ (Fold Change): 0.2. **(E)** Violin plots of TAM receptors and cathepsin D gene expression in three macrophage subtypes defined in B. **(F)** Representative confocal images of Axl^+^ and Axl^-^ macrophages in E11.5 rat > mouse chimeras. **(G)** Whole embryo chimerism analysis of E11.5 rat > mouse chimeras. Mouse embryos with four different genotypes are used for chimera generation. Each dot represents one chimeric embryo. The mean chimerism of each group is plotted as a black bar. Chimerism was quantified by flow cytometry. n = 17 (WT), 9 (Axl KO), 4 (Mertk KO), 8 (Axl, Mertk double KO). Statistical significance was assessed using unpaired two tailed t-test. n.s., not statistically significant; *p < 0.05, **p < 0.01, ***p < 0.001. See also Figure S3.

A total of 3065 cells were individually sorted and sequenced, with excellent gene count (median 7567) and read count (median 5.97×105). The species assignment for sorted cells were verified by cross-species genome mapping and cluster analysis (Figure S3A). *EGFP* transcript correlated well with EGFP fluorescence intensity as measured by index sort (Figure S3B). Rat-derived *EGFP* transcripts were detected most strongly in the rat cells, as expected, but were also detected in the eating macrophages, consistent with them having recently phagocytized a rat cell (Figure S3C). Macrophages isolated from different embryonic stages clearly separated on the UMAP plot, highlighting the maturation of macrophages from E9.5 to E11.5 (Figure S3D). Unsupervised clustering yielded distinct pre-macrophage and macrophage populations, but did not separate eating versus non-eating macrophages (Figure 3C). Differential gene expression analysis of E11.5 macrophages showed that eating macrophages, compared to noneating macrophages, upregulated genes involved in protein catabolism (Ctsd, Ctsh, Ctsz, Lgmn) and macrophage inflammation (Sparc, Spp1), indicating an activated phenotype. Notably, eating macrophages selectively upregulated Axl, a member of the TAM receptor family which plays crucial role in phagocytosis^51^ (Figure 3D). Tyro3 and Mertk (other two members of the TAM receptor family) were not upregulated (Figure 3E). Classical markers of M1/M2 polarization were not differentially expressed at either the RNA or protein level between eating vs noneating macrophages (Figure S3E). Overall, these results show that eating macrophages are transcriptomically distinct from their resting counterparts.

To investigate whether Axl expression marks macrophages involved in xenogeneic cell recognition, we performed immunostaining on E11.5 rat > mouse chimeras. A subset of F4/80+ macrophages exhibiting strong cell-surface expression of Axl were frequently found contacting with xenogeneic rat cells (Figure 3F). In contrast, macrophages lacking Axl expression were rarely observed in close proximity to rat donor cells. These observations suggest that Axl expression on macrophages may play an important role in mediating recognition and interaction with xenogeneic donor cells. To test whether Axl plays a functional role in macrophage-mediated xenophagocytosis, we first established an *in vitro* assay to recapitulate this phenomenon. Embryoid bodies (EBs) were generated from EGFP-labeled mouse or rat ESCs to mimic early embryonic development. Day 10 EBs were dissociated into single cells and co-cultured with mouse bone marrow-derived macrophages (Figure S3F). Flow cytometry analysis revealed that mouse macrophages exhibited approximately three-fold higher phagocytic activity against rat cells compared to mouse cells. Addition of BMS777607, a pan-inhibitor of the TAM receptor family^52^, significantly reduced xenophagocytosis frequency (Figure S3F).

Given that macrophages from E11.5 chimeras expressed high levels of both Axl and Mertk but minimal levels of Tyro3 (Figure 3E), we next sought to determine whether Axl and Mertk have redundant or distinct roles in xenophagocytosis. Immunostaining confirmed co-expression of Mertk and Axl in Axl^+^F4/80^+^ macrophages (Figure S3G). To directly test the functional contributions of these receptors, we generated single and double knockout embryos for Axl and Mertk by electroporating sgRNA/Cas9 RNP complex into mouse zygotes^44^, followed by injection of rat PSCs to produce rat > mouse chimeras. Chimerism analysis at E11.5 revealed significantly increased rat donor chimerism in Axl knockout embryos compared to wild-type controls, whereas Mertk knockout embryos showed no significant increase (Figure 3G). Double knockout of Axl and Mertk resulted in slightly higher chimerism compared to Axl single knockout embryos, suggesting a potential additive effect between these two receptors. Taken together, these results indicate that Axl is the predominant receptor mediating macrophage recognition and phagocytosis of xenogeneic donor cells, with Mertk playing a secondary, supportive role.

### Xenogeneic donor cells in chimeras display elevated phosphatidylserine

Given that macrophage recognition of xenogeneic donor cells was mediated predominantly through the receptor Axl, we next sought to identify the ligand responsible for triggering this interaction. Axl is known to recognize phosphatidylserine (PtdSer)^51^, a phospholipid normally restricted to the inner leaflet of the plasma membrane^53^ but exposed on the cell surface during apoptosis, serving as a canonical “eat-me” signal for macrophages^54,55^.

To determine whether PtdSer exposure was elevated on xenogeneic donor cells, we performed Annexin-V staining^56^ followed by flow cytometry analysis specifically on live (propidium iodide-negative) cells isolated from chimeric embryos. Donor and host cells exhibited comparable levels of surface PtdSer in intraspecies chimeras (mouse > mouse or rat > rat) (Figures 4A and 4B). In contrast, xenogeneic donor cells consistently displayed significantly higher levels of surface PtdSer compared to the host cells in interspecies chimeras (rat > mouse or mouse > rat) (Figures 4A and 4B). These results suggest that the xenogeneic developmental environment specifically induces elevated exposure of the “eat-me” signal PtdSer on donor cells. Traditionally, surface exposure of PtdSer is considered a hallmark of early apoptosis^55^. However, our earlier analyses clearly demonstrated that xenogeneic donor cells were not apoptotic. Therefore, we hypothesized that elevated PtdSer exposure on xenogeneic donor cells might disrupt the balance between “eat-me” and “don’t eat me” signals, leading to their selective phagocytosis by host macrophages independently of apoptosis.

**Figure 4.**
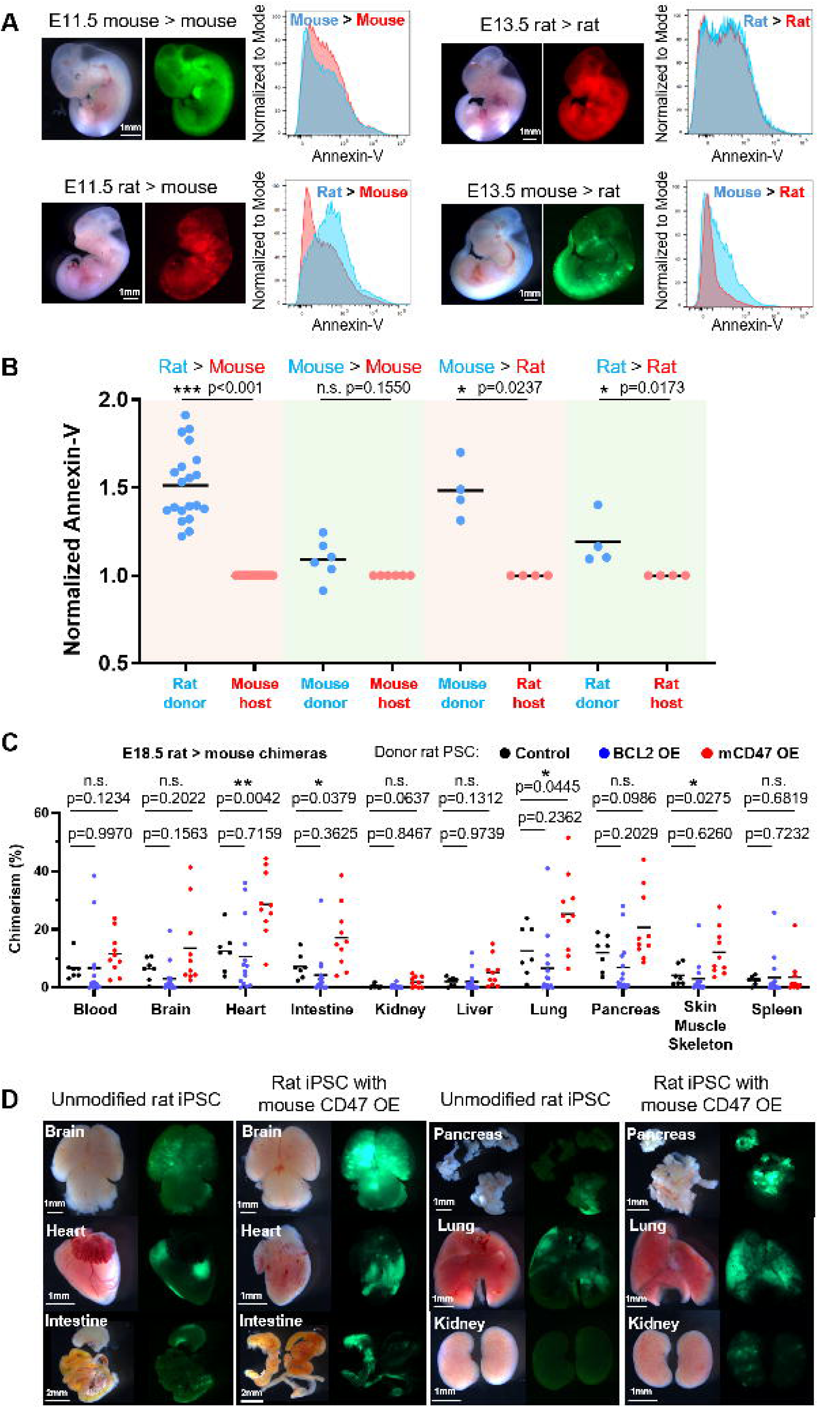
Xenogeneic donor cells exhibit elevated phosphatidylserine exposure and mouse CD47 overexpression on donor rat cells increases interspecies chimerism. **(A)** Representative flow cytometry results of Annexin-V staining in E11.5 mouse□>□mouse, E13.5 rat□>□rat (intraspecies) and E11.5 rat□>□mouse, E13.5 mouse□>□rat (interspecies) chimeras. Histograms show Annexin-V signal for donor (blue) and host (red) cells. **(B)** Quantification of donor-cell Annexin-V signal, normalized to host-cell signal for each chimera. Each dot represents one chimeric embryo. The mean Annexin-V of each group is plotted. n = 20 (rat□>□mouse), 6 (mouse□>□mouse), 4 (mouse□>□rat), and 4 (rat□>□rat). Statistical significance was assessed using paired two tailed t-test. n.s., not statistically significant; *p < 0.05, **p < 0.01, ***p < 0.001. **(C)** Donor chimerism in ten organs from E18.5 rat□>□mouse chimeras generated with rat iPSCs overexpressing EGFP (control), rat Bcl2, or mouse CD47. Each dot represents one organ dissected from one chimera. Black bars indicate group means. Chimerism was quantified by ddPCR. n = 7 (control), 14 (rat Bcl2 OE), and 10 (mouse CD47 OE). Statistical significance was assessed using unpaired two tailed t-test. n.s., not statistically significant; *p < 0.05, **p < 0.01. **(D)** Representative bright-field and fluorescence images of organs from E18.5 rat > mouse chimeras generated with control or mouse CD47-overexpressing rat iPSCs. EGFP indicates donor cell contribution. See also Figure S4.

To test this hypothesis, we designed two strategies aiming at suppressing xenophagocytosis: (1) overexpression of rat Bcl2 in rat PSCs to inhibit apoptosis^11,57^, and (2) overexpression of mouse Cd47, a major “don’t eat me” signal^58–60^, in rat PSCs to counterbalance elevated PtdSer exposure. Rat PSCs with overexpression of Bcl2 or mouse CD47 were generated and validated at the protein level (Figures S4A and S4B). These engineered rat PSCs were injected into wild-type mouse blastocysts to make rat > mouse chimeras. Chimerism analysis of organs at E18.5 revealed that rat Bcl2 overexpression did not significantly increase donor chimerism compared to controls (Figure 4C). In contrast, overexpression of mouse Cd47 significantly enhanced donor chimerism in multiple organs, including heart, intestine, lung and skin/muscle (Figures 4C and 4D). These results further support our conclusion that macrophage-mediated xenophagocytosis occurs through an apoptosis-independent mechanism driven by elevated PtdSer exposure on viable donor cells.

### PU.1 knockout enhances human cell chimerism in mouse embryo and yolk sac

Human PSCs typically exhibit minimal contribution to mouse embryos^13,14,16,17,19,61^, likely due to significant evolutionary divergence between these two species. We next investigated whether xenophagocytosis also affects human cell survival in human > mouse chimeras. To test this, we utilized an engineered human iPSC line maintained in the primed pluripotent state^62^, in which human BCL2 was constitutively overexpressed to inhibit apoptosis, and human MYC was expressed under a tetracycline-inducible promoter to promote proliferation (Figure 5A). These genetic modifications were found to be essential for achieving detectable human cell engraftment in mouse embryos past E9.5^62^. EGFP labeled human iPSCs were injected into mouse blastocysts to generate human > mouse chimeras. Doxycycline was supplied to the foster mother until E9.5 before macrophages completed colonization. Human cells were readily identified in E9.5 mouse embryos (Figure S5A). A decline in chimerism was observed at E11.5 (mean chimerism: 0.24% at E9.5 and 0.02% at E11.5) (Figures S5B and S5C), similar to what we observed in rat > mouse chimeras (Figure 1A). Although chimerism was limited in the embryo proper, human cells were frequently observed in the heart (Figures 5B and S5B). After careful analysis of chimeras at E11.5 including extraembryonic tissues, we identified human cells in the yolk sac with higher frequency and chimerism compared to embryo proper (Figures 5B and 5C). Flow cytometry analysis identified CD11b^+^, F4/80^+^, EGFP^+^ macrophages in yolk sac at E11.5, suggesting active xenophagocytosis in human > mouse chimeras (Figure 5C). Annexin-V staining revealed that the xenogeneic human donor cells displayed higher levels of surface PtdSer compared to the mouse host cells in the chimeric yolk sac samples (Figure S5D).

**Figure 5.**
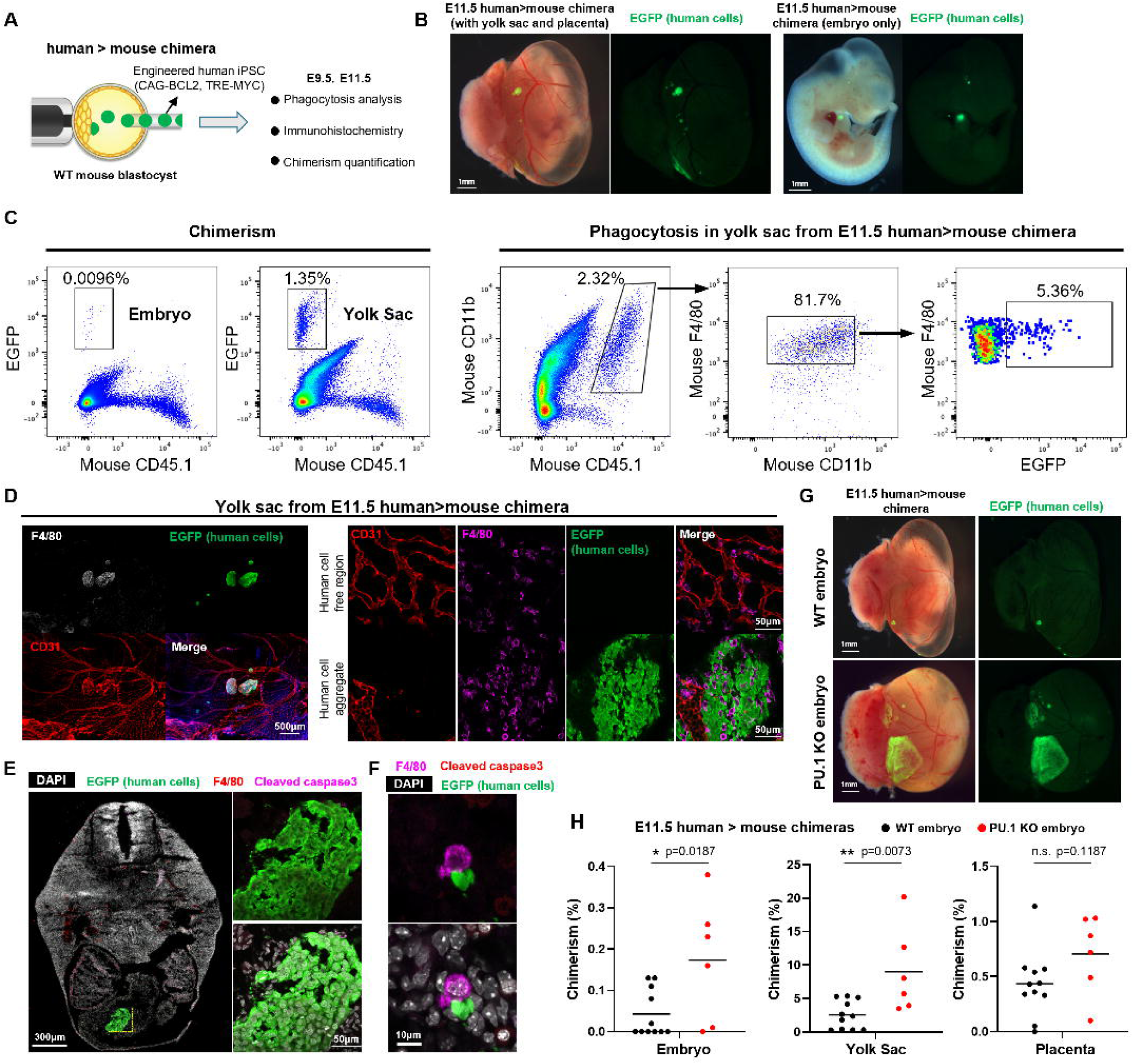
Macrophage deficiency in host embryos enhances human donor cell chimerism in extraembryonic and embryonic tissues. **(A)** Schematic of chimera generation by injecting genetically engineered human iPSCs into WT mouse blastocysts. Chimeras were analyzed at E9.5 and E11.5. Donor chimerism was quantified by ddPCR, spatial distribution was assessed by immunostaining, and phagocytosis was analyzed by flow cytometry. **(B)** Bright-field and fluorescence images of an E11.5 human□>□mouse chimera before (left) and after (right) removal of yolk sac and placenta. EGFP indicates human donor cells. **(C)** Flow cytometry analysis of embryo and yolk sac from an E11.5 human□>□mouse chimera. The human donor chimerism in the embryo proper and yolk sac are shown. A subset of CD45.1□CD11b□F4/80□ mouse macrophages containing EGFP□ signal were classified as “eating macrophages”. **(D)** Left: fluorescence images of E11.5 yolk sac tissue stained for EGFP (human cells), CD31 (endothelial cells), and F4/80 (mouse macrophages). Right: magnified views of regions with and without human donor cells. **(E)** Transverse section of an E11.5 human□>□mouse chimera stained for EGFP (human cells), F4/80 (mouse macrophages), and cleaved caspase-3 (apoptotic marker). A cluster of EGFP□ human cells lacking cleaved caspase-3 staining is observed within the embryonic heart. Right: magnified view of a human cell outside the heart in contact with a F4/80□ macrophage. **(F)** Magnified view of EGFP^+^ human cells outside the heart in contact with a F4/80□ mouse macrophage. **(G)** Bright-field and fluorescence images of E11.5 human□>□mouse chimeras before removal of yolk sac and placenta. Chimeras were generated using WT or PU.1 knockout host embryos. **(H)** Human donor chimerism in embryo proper, yolk sac and placenta of E11.5 human□>□mouse chimeras. Chimeras were generated using WT or PU.1 knockout mouse embryos. Each dot represents tissue from one chimeric embryo. Black bars indicate mean chimerism of each group. Chimerism was quantified by ddPCR. n = 11 (WT), 6 (PU.1 KO). Statistical significance was assessed using unpaired two tailed t-test. n.s., not statistically significant; *p < 0.05, **p < 0.01. See also Figures S5 and S6.

Immunostaining of E11.5 chimeric yolk sacs revealed abundant and evenly distributed macrophages at this developmental stage (Figure S5E). Notably, macrophages accumulated at higher densities in regions containing human donor cells (Figure 5D), confirming active xenophagocytosis. Aggregates of EGFP^+^, cleaved caspase-3-negative human cells were observed within cardiac chambers, indicating that these human cells were viable and not apoptotic (Figure 5E). Interestingly, our previous experiment showed that the heart was largely devoid of F4/80^+^ macrophages at this stage (Figures 1F and S1D), potentially providing a favorable environment for human cell survival. Although chimerism outside the heart remained extremely low, we still observed mouse macrophages in direct contact with viable (cleaved caspase-3-negative) human cells (Figure 5F).

To directly test whether reducing xenophagocytosis could enhance human donor cell chimerism, we injected hiPSCs into PU.1 deficient mouse blastocysts and compared with control embryos. Next, we quantified human donor chimerism using a highly sensitive ddPCR assay at E11.5. Given the unique propensity of human cells to contribute preferentially to mouse extraembryonic tissues, we dissected embryo proper, yolk sac and placenta from each chimera and analyzed chimerism separately. PU.1-deficient embryos exhibited significantly increased human chimerism in embryo proper and yolk sac compared to controls, with yolk sac having the most prominent increase in chimerism (up to 20%) (Figures 5G and 5H). The absolute percentage of chimerism in the embryo proper remained low (up to 0.4%) in PU.1 deficient embryos, although clearly outperformed the controls. The extremely low chimerism at E9.5 (Figures S5A and S5C) suggests the existence of other xenogeneic barriers that are unique to human > mouse chimeras, which happen before the emergence of macrophages. Nevertheless, these data indicate that a xenophagocytosis barrier also applies to human > mouse chimeras.

## DISCUSSION

Here we discovered that xenophagocytosis is a major barrier preventing the generation of high-chimerism interspecies chimeras between mouse and rat. Furthermore, our data strongly suggests that mouse macrophages also attack human cells in human > mouse chimeras, supporting the generalizability of xenophagocytosis across different species. Based on our findings, we propose the following model (Figure 6A): In early rat > mouse chimeras, xenogeneic rat donor cells display elevated levels of phosphatidylserine (PtdSer) on their cell surface through an apoptosis-independent mechanism. Although PtdSer exposure is classically associated with apoptosis^54^, it is also known to occur in viable cells under certain physiological or stress conditions^63–65^. This elevated PtdSer acts as an “eat-me” signal recognized by host macrophages through TAM receptors, particularly Axl. The imbalance between elevated “eat-me” signals and insufficient “don’t eat me” signals sensitizes xenogeneic donor cells as targets of the host macrophages. Mouse macrophages colonize embryos starting around E9.5 and mature by E11.5, at which point Axl+ macrophages actively recognize and eliminate xenogeneic rat cells. This selective removal of viable, PtdSer-positive donor cells aligns with the concept of phagoptosis—cell death initiated by phagocytic uptake rather than intrinsic apoptosis. Supporting this model, we have shown that genetic ablation of host macrophages significantly increases xenogeneic chimerism. Similarly, overexpression of the species-matched “don’t eat me” signal mouse Cd47 on rat donor cells partially restores the balance between “eat-me” and “don’t eat me” signals, protecting rat cells from xenogeneic attack.

**Figure 6.**
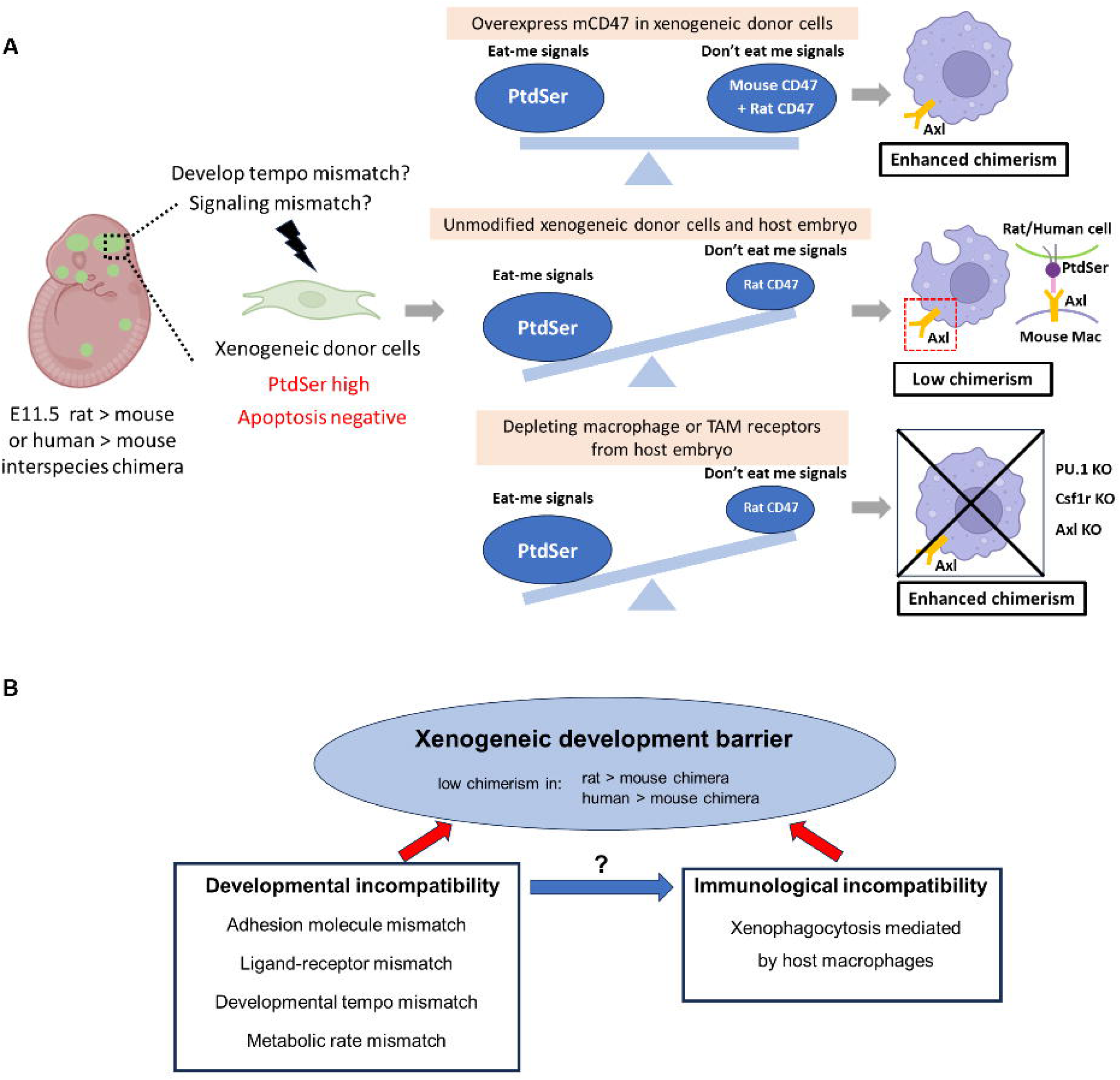
Schematic model of xenogeneic phagocytosis as a component of the barrier to xenogeneic organogenesis. **(A)** Proposed mechanism in early-stage rat□>□mouse chimeras. Elevated phosphatidylserine (PtdSer) exposure on non-apoptotic donor rat cells serves as an “eat-me” signal recognized by Axl on host macrophages. The imbalance between “eat-me” and “don’t eat-me” signals renders donor cells susceptible to phagocytosis. Xenogeneic clearance can be mitigated by either depleting host macrophages or overexpressing mouse CD47 on donor rat cells. **(B)** Conceptual integration of developmental and immunological incompatibility as components of the xenogeneic organogenesis barrier. Macrophages are identified as key effectors in eliminating xenogeneic donor cells. Developmental stress arising from interspecies mismatch may contribute to donor cell susceptibility.

Previous studies of the xenogeneic barrier during embryonic development are primarily from a developmental biology perspective^11–17,19,45^. Because long-term coexistence of donor and host cells was readily achieved in syngeneic and allogeneic intraspecies chimeras generated at the blastocyst stage, early studies tacitly applied the same assumption of “built-in immune tolerance” to interspecies chimeras. Our findings challenge this assumption, demonstrating that immunological incompatibility mediated by innate immune cells, specifically macrophages, is a critical component of the xenogeneic barrier even before the emergence of adaptive immunity. The developmental and immunological incompatibilities may be intrinsically connected (Figure 6B). Multiple developmental mismatches (e.g., signaling factor incompatibility, adhesion molecule mismatch, differences in developmental tempo) could induce cellular stress in xenogeneic donor cells, potentially leading to elevated PtdSer exposure. Future studies will aim to dissect the precise extrinsic factors responsible for inducing PtdSer exposure in xenogeneic donor cells.

Although we successfully increased chimerism by overexpressing mouse Cd47, the effect was not as robust as PU.1 knockout mediated macrophage ablation. This may reflect the diversity and tissue-specificity of “don’t eat me” signals expressed in different organs. Indeed, recent studies in cancer immunology have identified multiple “don’t eat me” signals, including Beta-2-microglobulin^66^, Cd24^67^, and PD-1^68^, each interacting with distinct macrophage receptors. Tissue-specific expression patterns of these inhibitory signals may explain the variable effectiveness of Cd47 overexpression across different organs. Future studies testing combinations of these signals may further increase interspecies chimerism.

Generating human organs in mouse embryos remains challenging, as human cells contribute minimally to the embryo proper (approximately 1 in 2,500 cells in human > mouse chimeras at E11.5 as shown in Figure 5H). This limited contribution is largely due to early segregation of human cells into extraembryonic tissues before E6.5, well before the emergence of macrophages^62^. Nevertheless, our data clearly demonstrates macrophage-mediated attack on human cells in the yolk sac, highlighting xenophagocytosis as an important barrier to consider when generating human organs in livestock.

Recent pig-to-human xenotransplantation studies have marked a major step in transplantation medicine^69–72^. However, despite the heroic number of genomic edits made and intense immunosuppression, these porcine organs still failed within months after transplantation into human patients^70,73,74^, likely due to remaining interspecies incompatibilities. Single-cell and spatial analyses of gene-edited pig-to-human grafts nevertheless revealed a consistent influx of recipient macrophages into the transplanted porcine organs^73,75^. Their presence alone raises the possibility that phagocytic elimination of xenogeneic donor cells—akin to the xenophagocytosis we define in embryos—may also contribute to graft failure. Our findings therefore complement current xenotransplantation strategies and suggest that macrophage-centered “eat-me / don’t eat me” checkpoints deserve closer investigation.

An additional important observation from our study was the clear difference in nuclear DNA staining patterns between mouse and human cells. We found that DAPI staining consistently produced a uniform nuclear staining pattern in human cells, whereas mouse nuclei exhibited bright, punctate DNA speckles independent of tissue type (Figures S6A and S6B). This species-specific nuclear staining pattern was also observed in several human-mouse xenotransplantation studies^76–80^. It provides a reliable and convenient criterion for distinguishing human donor cells from mouse host cells in chimeric embryos, complementing traditional fluorescent protein labeling approaches. Historically, many researchers have relied primarily on fluorescent protein markers (e.g., EGFP) to detect and quantify human donor chimerism in human > mouse chimeras. However, fluorescent protein signals can be confounded by tissue autofluorescence and can also be detected in cell debris, leading to potential misinterpretation and overestimation of donor cell contribution. Therefore, previous reports showing human > mouse chimeras need to be reevaluated based on this simple but reliable DAPI staining pattern. We strongly recommend that future chimera studies incorporate more rigorous and complementary methods, such as highly sensitive ddPCR assays and careful examination of nuclear staining patterns, to accurately validate and quantify donor cell engraftment. Such approaches will ensure robust and reproducible assessment of interspecies chimerism, particularly when evaluating human cell contributions in mouse host embryos.

By bridging developmental biology and immunology, our findings uncover xenophagocytosis as a key innate barrier to interspecies chimerism. We propose strategies, such as genetic engineering of host embryos or donor cells, to overcome this xenogeneic barrier. These insights provide a foundation for improving efforts to generate human organs in livestock. Regardless of specifics, a fundamental question remains: It is difficult to believe that this mechanism exists solely for the preservation of the species. What role might it have played in the normal processes of animal evolution and development? Clarifying the purpose of this mechanism remains a highly interesting question for future studies.

## STAR Methods

### ESCs and iPSCs culture

In this study, all mouse and rat ESC and iPSC lines were maintained on mitomycin-C-treated CF1 mouse embryonic fibroblast (MEF) feeder cells made in house. Undifferentiated mouse ESCs (SGE2 line, XY) were maintained in DMEM medium (Gibco, 11960044) containing 10% ESC qualified FBS (Sigma-Aldrich, ES009-M), 1000 units/mL mouse Lif (Sigma-Aldrich, ESG1106), 1μM PD0325901 (Selleckchem, S1036), 3 μM CHIR99021 (Selleckchem, S2924). Undifferentiated rat iPSCs (T1-3 line, XY) were maintained in N2B27 basal medium supplemented with 1000 units/mL rat Lif (Sigma-Aldrich, ESG2206), 1 μM PD0325901, 1 μM CHIR99021. Undifferentiated rat ESCs (Wistar-tdt-rESC-No7 line, XY) were maintained in N2B27 basal media supplemented 1000 units/mL rat Lif (Sigma-Aldrich, ESG2206), 1 μM PD0325901, 3 μM CHIR99021. The SGE2 mESC and T1-3 rat iPSC lines were established and validated from our previous studies^7,81^. The Wistar-tdt-rESC-No7 line was derived in this study.

Engineered human iPSCs (cell line PB004#1 with constitutive overexpression of BCL2 and doxycycline inducible MYC) were maintained on iMatrix-511 (Nacalai USA, 892021) coated 12 well plate and cultured in StemFit basic 04 complete medium (Amsbio, SFB-504-CT).

### Genetic modification of human iPSC

The human iPSC line PB001#4 was established from peripheral blood cells and reprogrammed using Sendai virus vector expressing *OCT4*, *SOX2*, *KLF4,* and *MYC*. The collection of peripheral blood samples and iPSC generation were approved by the ethical committee of the University of Tokyo.

A human *BCL2*-expressing lentiviral vector and a Venus-expressing lentiviral vector were constructed based on the CS-CAG-GFP lentiviral vector. The Dox-inducible human MYC-expressing lentiviral vector was derived from the lentiviral vector CS-TRE-PRE-Ubc-rtTA-IRES2-EGFP^82^. The open reading frames (ORFs) of BCL2 and MYC were amplified from cDNA synthesized from total RNA extracted from the human iPSC line PB001. The BCL2 ORF was inserted downstream of the CAG instead of GFP, and a puromycin resistance gene was inserted downstream of the IRES sequence. An rtTA-T2A-Venus was inserted downstream of the CAG promoter in place of GFP. The *MYC* ORF was inserted downstream of the TRE, followed by a Zeocin resistance gene downstream of IRES2, using the In-Fusion Snap Assembly Master Mix.

Lentiviruses were produced by co-transfection of the lentiviral expression vector plasmid with packaging plasmids (pMDL-gag/pol-PRE and pCMV-VSV-G-Rev) into 293T cells. Cells were seeded at 3.5×10^6^ cells per 10 cm dish one day before transfection. Transfections were carried out using the calcium phosphate method with 15 µg of the lentiviral vector and 10 µg each of the packaging plasmids. The medium was replaced 16 h after transfection, and the cells were incubated for 48 h. Lentiviruses were concentrated by ultracentrifugation at 4000 x *g* for 2 h, and the pellet was resuspended in 1/250th volume of PBS. Concentrated viruses were stored at –80°C.

GFP-, Venus-, *BCL2*- and *MYC*-lentivirus were introduced into feeder-free hPSCs by adding them to the culture medium for 1-2 days. Puromycin (500 ng/mL) or zeocin (5 µg/mL)-resistant clones were selected after 3 days of lentivirus transduction. Venus- or GFP-positive hiPSC cells were purified by FACS sorting and subsequently propagated.

### Derivation of rat ESC line

We generated a transgenic Wistar rat line that had tdTomato driven by the constitutive active CAG promoter. Wild-type wistar female rats were mated with tdTomato expressing male rats. Rat blastocysts were flushed from time-pregnant rats’ uteruses at E4.5. Zona pellucida was removed by placing rat blastocysts in acid Tyrode solution (CytoSpring, AT001). Each zona pellucida free rat blastocyst was transferred into one well in the 12 well plate with already plated MEF feeders at high density (400K MEFs in each well). The derivation medium was the same as culture medium for Wistar-tdt-rESC-No7 line. After 7 days, blastocysts with clear and round outgrowth were picked for single cell dissociation. The single cells were replated in a new well of 12 well plate with dense MEF feeders (400K MEFs in a single well). Among all lines established, line No.7 inherited the CAG-tdTomato transgene, had XY genotype, and was able to generate high chimerism rat>rat chimeras with high frequency. Therefore, this line (Wistar-tdt-rESC-No7) was used for the current study.

### Mouse embryo collection and culture

3-4-week-old CD1 females were super ovulated by injecting 7.5 units of Pregnant Mare Serum Gonadotropin (PMSG, Lee BioSolutions, 493-10-10) intraperitoneally. 48 hours later, 7.5 units of Human Chorionic gonadotropin (hCG, Sigma-Aldrich, CG10-1VL) was injected and the females were set up mating with CD1 stud males. Females with plug were identified next day and noon was defined as embryonic day 0.5 (E0.5). Morula stage embryos were flushed from the oviduct and first half of uteruses at E2.5. Morulas were cultured in KSOM-AA medium (Sigma-Aldrich, MR-106-D) overnight to develop to blastocysts. Blastocysts with well-developed cavities were used for microinjection.

### Rat embryo collection and culture

7-10 weeks Wistar female rats were synchronized for their estrus cycles by intraperitoneally injection of 40μg luteinizing hormone-releasing hormone (LH-RH, Sigma-Aldrich, L4513)^83,84^. 96 hours later, females were set up mating with Wistar stud males. Females with plug were identified next day and noon was defined as embryonic day 0.5 (E0.5). Rat blastocysts were flushed from the uteruses at E4.5. Rat blastocysts were cultured in mR1ECM medium (Cosmo Bio, CSR-R-M191) for 2-4 hours. Blastocysts with well expanded cavities were used for microinjection.

### Genome editing of mouse embryos by Cas9/gRNA ribonucleoprotein (RNP) electroporation

CD1 female mice were super ovulated and mated with stud males as mentioned above. Zygotes were collected from the swollen ampulla of the female mice at E0.5. Cumulus cells were removed by incubating the cumulus-oocyte complex in the M2 embryo media (CytoSpring, M2103) containing 0.3 mg/ml hyaluronidase (Sigma-Aldrich, H4272). The cumulus cell free zygotes were cultured in KSOM-AA medium for 2-4 hours before electroporation. Before electroporation, the zygotes with clear two pronucleus were selected and washed three times in M2 medium. 20-30 zygotes were transferred into the small space between two electrodes of the BEX CFB16-HB electroporator (BEX CO.,LTD). The space between electrodes was pre-filled with 5uL Opti-MEM medium (Gibco, 31985062) containing 100ng/μL S.p. HiFi Cas9 Nuclease V3 protein (Integrated DNA Technologies, 1081061) and 100ng/μL sgRNA (Synthego). The electroporation program for CD1 zygotes was: 25 V, 3ms ON, 97 ms OFF, Pd Alt 3 times^44,45^. After electroporation, zygotes were washed three times with M2 medium and transferred to KSOM-AA medium for culture for 4 days. The KSOM-AA medium was changed every other day. 4 days after culture, embryos that developed into blastocyst stage were selected for either direct embryo transfer or microinjection.

The sgRNA targets used in this study were:

Mouse Spi1: 5’-tgataagggaagcacatccg - 3’

Mouse Csf1r: 5’-cgcagggtcaccgtttcacc- 3’

Mouse Axl: 5’-tcgaagccacaccacctcag- 3’

Mouse Mertk: 5’-ttgatgctgtggaggtcacc- 3’

### Generation of PU.1 and Csf1r transgenic mouse lines

Wild-type CD1 mouse zygotes were electroporated with Cas9/sgRNA RNP that contained sgRNA targeting either mouse Spi1 (5’-tgataagggaagcacatccg - 3’) or Csf1r (5’-cgcagggtcaccgtttcacc- 3’). Zygotes were cultured in KSOM-AA medium until reaching the blastocyst stage before embryo transfer. 15-20 well developed blastocysts were transferred into uteri of pseudo pregnant recipient CD1 female mice (2.5 days post coitum). Male mice that survived to 8 weeks were genotyped and screened for heterozygous founders. The founder males were crossed with wild-type CD1 females for germline transmission of the gene edited allele. Germline transmitted F1 mice with heterozygous allele for PU.1 or Csf1r were identified and further backcrossed with wild-type CD1 for three generations to establish a stable colony. Eventually, mice that are heterozygous for PU.1 (−1bp/+) and Csf1r (−34bp/+) were phenotypically validated and maintained in the lab.

### Chimera generation by embryo manipulation

In this study, all intraspecies and interspecies chimeras were generated by injecting ESCs or iPSCs into blastocysts stage embryos. A laser-based micromanipulator (RI Saturn 5 Active™ laser system, CooperSurgical, Inc.) was used to break the zona pellucida and the trophectoderm and single ESCs or iPSCs were injected into blastocysts cavities and placed near the inner cell mass. More specifically, the details for different chimera conditions were described below.

For generating mouse>mouse or rat>mouse chimeras, mouse ESCs or rat ESC/iPSCs were dissociated into single cells by Tryple Express Enzyme (Thermo Scientific, 12604021) and resuspended in corresponding ESC or iPSC culture medium. 6-8 mouse ESCs or 5-6 rat ESCs/iPSCs were injected into each mouse blastocyst. After injection, the chimeric embryos were cultured for 1-2 hours in KSOM-AA medium for recovery. Chimeric mouse blastocysts were transferred into the uterus of pseudo pregnant CD1 female mice (2.5 days post coitum).

For generating rat>rat or mouse>rat chimeras, rat ESC/iPSCs or mouse ESCs were dissociated into single cells by Tryple Express Enzyme and resuspended in corresponding ESC or iPSC culture medium. 6-8 rat ESCs/iPSCs or 5-6 mouse ESCs were injected into each rat blastocyst. After injection, the chimeric embryos were cultured for 1-2 hours in mR1ECM medium for recovery. After recovery, the rat blastocysts were transferred into the uterus of pseudo pregnant Wistar female rats (3.5 days post coitum).

For generating human>mouse chimeras, human iPSCs were dissociated into single cells using Accutase (STEMCELL Technologies, 07920) and resuspended in StemFit Basic 04 complete medium supplemented with 5μM ROCK Inhibitor Y-27632 (STEMCELL Technologies, 72307). 4-5 human iPSCs were injected into each mouse blastocyst. After injection, the chimeric embryos were cultured for 1-2 hours in KSOM-AA medium supplemented with 2.5μM Y-27632. After recovery, the chimeric mouse blastocysts were transferred into the uterus of pseudo pregnant CD1 female mice (2.5 days post coitum). To activate the inducible MYC expression in human cells, doxycycline containing drinking water consisting of 2mg/mL doxycycline hyclate (Fisher Scientific, AAJ6057922) and 5% sucrose (Sigma-Aldrich, S0389) was given to pseudo pregnant mice from E2.5 to E9.5.

### Immunohistochemistry and microscopy

E11.5 rat>mouse chimeras, E13.5 mouse>rat chimeras and dissected organs from E18.5 rat>mouse chimeras were fixed in 4% paraformaldehyde (Santa Cruz Biotechnology, SC-281692) at 4 °C overnight. Fixed tissues were washed 3 times with PBS before transferring into 30% sucrose at 4 °C overnight. Tissues were then embedded in Tissue Plus O.C.T. compound (Fisher Scientific, 23-730-571) and sectioned at 15 μm to obtain frozen sections. For staining, frozen sections were thawed to room temperature and rehydrated in PBS. Sections were permeabilized ad blocked with 5% normal donkey serum (SouthernBiotech, 0030-01) in PBS supplemented with 0.3% Triton-X100 (Sigma-Aldrich, T9284) for 1 hour at room temperature. Primary antibodies were diluted in the blocking buffer and applied on sections for overnight incubation. Slides were washed three times with PBS before being incubated with secondary antibodies and DAPI (1ug/mL, Thermo Fisher Scientific, 62248) in blocking buffer for 1 hour at room temperature. Slides were washed three times with PBS before being mounted using Fluoromount-G (SouthernBiotech, 0100-01). Images were captured using either Keyence All-in-One fluorescence microscope BZ-X710 or a LSM980plus confocal microscope. A list of primary antibodies with dilution ratios can be found in Table S1.

### Embryonic body generation from mouse ESCs or rat ESCs/iPSCs

Mouse ESCs or rat ESC/iPSCs were dissociated into single cells by TrypLE and resuspended in ESC culture medium. MEF feeders were depleted by plating single cells on gelatin (Millipore Sigma, SF008) coated plate for half hour at 37°C. AggreWell 800 24-well plate (STEMCELL Technologies, 34850) was used to generate embryonic bodies (EBs). 1000 single cells were plated in each microwell and incubated overnight. Next day, embryonic bodies were transferred into ultra-low attachment 6-well plates (Corning, 3471) for long-term culture. DMEM with 10% FBS was used as culture medium for EBs to promote spontaneous differentiation. EBs were cultured for 10 days in vitro. Day 10 EBs were dissociated into single cells by incubating with Accutase for 1 hour at 37°C. Single cells from EBs were used for co-culture experiments with macrophages.

### Single cell dissociation of dissected organs for flow cytometry analysis

Organs (brain, heart, lung) from E18.5 mouse embryos or 2 weeks mice were dissected in ice cold PBS (Corning, 21-040-CV). Each organ was dissociated into single cells by incubating in a collagenase-based dissociation buffer^85^ for 30 minutes (heart) or 45 minutes (lung) at 37°C with constant shaking. The dissociation buffer consisted of collagenase IV (2.2mg/mL, Thermo Scientific, 17104019), BSA (0.1%, Sigma-Aldrich, A3294), DNase (125units/mL, Worthington Biochemical, LS002006), HEPES (20mM, Gibco, 15630106), CaCl_2_ (1mM, Sigma-Aldrich, 21115-100ML), Pluronic F-68 (1:100, Thermo Scientific, 24040032) in basal medium 199 (Sigma-Aldrich, M4530). The single cell suspension was filtered through a 100μm strainer (Fisher Scientific, 07-201-432) and pelleted down. The cell pellet was resuspended in FACS buffer (PBS with 2% FBS plus 0.1% Pluronic F-68) for flow cytometry analysis.

### Single cell dissociation of whole embryo for flow cytometry analysis

For E9.5-E11.5 mouse embryos and chimeras, E11.5-E13.5 rat embryos and chimeras, we used a “cold digestion” protocol^86^ to maximize the cell viability after single cell dissociation. Embryos were dissected in a Petri dish filled with ice-cold HBSS (Sigma-Aldrich, 55037C-1000ML). A papain based Neural Tissue Dissociation kit (Miltenyi Biotec, 130-092-628) was used. All incubation steps were performed at 10°C in a pre-chilled refrigerated centrifuge. Each embryo was incubated with 500μL Mix1 for 15 minutes at 10°C. Mix 2 was added and embryos were further incubated for 20 minutes at 10°C. Embryos were pipetted with a P1000 pipette every 10 minutes until fully dissociated. Dissociated single cells were filtered through a 100μm strainer and spun down at 500 g for 5 min at 4°C. The supernatant was aspirated, and the cell pellet was washed twice by resuspending in 5mL FACS buffer (PBS with 2% FBS plus 0.1% Pluronic F-68). The cells were centrifuged again at 500g for 5min at 4°C. After the final wash, the cells were resuspended in FACS buffer for flow cytometry analysis or scRNA-seq with index sorting.

### Genetic modification of rat iPSCs

To overexpress rat BCL2 in rat iPSC T1-3 line, the piggyBac transposon system was used. Rat Bcl2-P2A-EGFP was cloned after a constitutive active CAG promoter in a bicistronic piggyBac plasmid. An Ubc promoter-puromycin selection cassette within the piggyBac construct facilitates selection of cells with stable integration of the transgene. Rat iPSCs were cultured in 6-well plate before transfection. 3μg piggyBac plasmid (pPBCAG-rat BCL2-P2A-EGFP_hUBC-Puro) and 1μg helper plasmid (pCAG-transposase) were transfected into rat iPSCs with Lipofectamine 3000 (Life Technologies, L3000-015) following vendor’s recommended protocol. Cells containing the integrated transgene were enriched by adding puromycin (1μg/mL, Invivogen, ANT-PR-1) in culture medium for 3 days. Single cell derived sub-clones were established from the puromycin selected population. Clones with stable overexpression of rat Bcl2 was identified through western blot and used for rat>mouse chimera generation.

To overexpress mouse Cd47 in rat iPSC T1-3 line, a lentivirus-based overexpression system was used. Mouse Cd47-IRES2-EGFP sequence was cloned after a CAG promoter in a CSII-based lentiviral vector. Lentivirus was obtained by transfecting lentiviral transfer plasmid (CSII_CAG-mCD47-IRES2-EGFP-WPRE), pMD2.G packaging plasmid (Addgene, #12259) and psPAX2 envelope plasmid (Addgene, #12260) into HEK 293T cells with Lipofectamine 3000. The viral supernatant was collected twice at 48 hours and 72 hours after transfection. The pooled viral supernatant was filtered through a 0.45mm PES filter and concentrated by ultracentrifugation. Rat iPSCs were transduced by lentivirus and allowed to expand for four days. Cells with overexpression of mouse Cd47 were identified by staining with Brilliant Violet 421 conjugated anti-mouse Cd47 antibody (BioLegend, 127527). Rat iPSCs with stable and high expression level of mouse Cd47 were established through three consecutive rounds of FACS sorting. Early passage of Cd47 overexpressed rat iPSCs were used for generating rat>mouse chimeras.

### Genotyping of non-chimeras

The tails from mouse embryos or adult mice were lysed in a homemade crude lysis buffer (15mM Tris, 100mM NaCL, 5mM EDTA, 0.1% SDS, pH 8.4) supplemented with 1mg/mL proteinase K (ApexBio, K1037). PCR strips contain tail samples and lysis buffer were incubated at 56 degrees for 1 hour followed by heat inactivation of proteinase K at 80 degrees for 10 minutes. The crude lysates were spun down at 3000 g for 5 minutes. The supernatant contained genomic DNA was used for genotyping.

SeqAmp DNA polymerase (Takara Bio, 638509) was used for performing genomic PCR for all samples in this study. The following primer sets were used for different genes.

Mouse Spi1: forward primer: CACTCACTTCCGTATTAAGCCC, reverse primer: TCATAGGCCTTCACACATTCAC.

Mouse Csf1r: forward primer: GGCTTGGCCTGCTTAGTACA, reverse primer: TGTTGATGCCAGGAGCTAGC.

Mouse Axl: forward primer: ACATGAGATGGAAGCCGGAC, reverse primer: GATTCCCAACCCAGGTGGAG.

Mouse Mertk: forward primer: AGCTCTTGACCCTCCTTCCT, reverse primer: TCCACCATGAAATGCCAGCT.

The PCR products were run on 1.5% agarose gel. The PCR amplicons with correct DNA size were cut out and subjected to gel extraction with QIAquick Gel Extraction Kit (Qiagen, 28706). The purified DNA was sequenced by Sanger sequencing for genotype determination.

### Genotyping of chimeras

For intraspecies (mouse>mouse) and interspecies chimeras (rat>mouse) at early stages (E9.5 to E11.5), the whole embryo was dissociated into single cells by the above mentioned “cold digestion” protocol. 20,000 – 50,000 single cells from the recipient mouse embryos were sorted out by FACS aria II by gating on GFP-, tdTomato-population. The sorted cells were spun down and the cell pellet was lysed by tissue lysis buffer. The following genotyping procedures were the same as mentioned in the “Genotyping of non-chimeras” section.

For intraspecies (mouse>mouse) and interspecies chimeras (rat>mouse) at E14.5 and E18.5, the spleen from each chimera was dissected and dissociated into single cells in a collagenase-based dissociation buffer (mentioned in previous section) for 20 minutes at 37°C with constant shaking. The cells were pelleted down and then subjected to Ammonium-Chloride-Potassium (ACK) lysis (Thermo Scientific, A1049201) to remove red blood cells. After the ACK lysis, the remaining cells were stained with APC conjugated anti-mouse Cd45 antibody (BioLegend, 103111) before FACS analysis. The splenocytes from the mouse recipient embryos were sorted out by FACS by gating on GFP-, tdTomato- and mouse CD45+ population. 30, 000 splenocytes were sorted out and lysed for genotyping of each chimera. The detailed genotyping procedures were the same as mentioned in the “Genotyping of non-chimeras” section.

### Digital droplet PCR (ddPCR) assay for chimerism analysis

For rat>mouse chimeras harvested at E9.5, whole embryo was lysed in a homemade crude lysis buffer (15mM Tris, 100mM NaCL, 5mM EDTA, 0.1% SDS, pH 8.4) supplemented with 200ug/mL proteinase K^31^. PCR strips contain embryos and lysis buffer were incubated at 56 degrees for 1 hour followed by heat inactivation of proteinase K at 75 degrees for 10 minutes. Crude lysates were homogenized by rigorous pipetting and spun down at 3000 g for 5 minutes. The supernatant containing the genomic DNA was used for ddPCR analysis.

For mouse>mouse, rat>mouse and human>mouse chimeras harvested at E11.5, E14.5 and E18.5, whole embryo (E11.5 or E14.5) or dissected organs from E18.5 chimeras were lysed in a stronger tissue lysis buffer (15mM Tris, 100mM NaCL, 5mM EDTA, 1% SDS, pH 8.4) supplemented with 1mg/mL proteinase K. Eppendorf tubes contain embryos or organs were incubated at 56 degrees for 8 hours with rigorous pipetting every two hours to fully homogenize the samples. After which the samples were heat inactivated at 75 degrees for 30 minutes. To precipitate SDS, 1M potassium chloride (KCL) solution was added into each sample with a 10% volume/volume ratio and the samples were put on ice for 10 minutes. After which the crude lysates were spun down at 3000 g for 5 minutes to allow SDS fully precipitate. The supernatant containing genomic DNA was used for ddPCR analysis.

For mouse>mouse chimeras, a ddPCR single nucleotide discrimination assay was used to count donor and host alleles as previously described^31,45^. Briefly, a primer pair was used to amplify the tyrosinase locus in the extracted DNA (forward, mTyr-F/1, 1.8 uM, AATAGGACCTGCCAGTGCTC; reverse, mTyr-R/1, 1.8 uM, TCAAGACTCGCTTCTCTGTACA). Two probes were added that bind to either the albino allele to detect the CD1 host embryo or the wildtype alleles to detect C57BL/6 donor cells (albino probe, mTyr-alb-P/1, 0.25 uM, fluorescein amidites (FAM)-cttaGagtttccgcagttgaaaccc-ZEN/IBQ; wildtype probe, mTyr-wt-P/1, 0.25 uM, hexachloro-fluorescein [HEX]-cttaCagtttccgcagttgaaaccc-ZEN/IBQ). The primers/probes and extracted DNA were mixed in a combined ddPCR reaction and run in standard conditions as described below. Percent chimerism of the donor cells was measured as the *wildtype* copies/ul divided by the sum of the *wildtype* and *albino* copies/ul.

For rat>mouse and mouse>rat chimeras, ddPCR primers and probes were designed to target either mouse P53 (forward, mP53-F/1, 0.9 uM GTGCTCACCCTGGCTAAAGT; reverse, mP53-R/1, 0.9 uM, AGGAGGATGAGGGCCTGAAT; probe, mP53-P/1, 0.25 uM, [HEX]-TGGGACCATCCTGGCTGTAGGT-ZEN/IBQ) or rat P53 (forward, rP53-F/1, 0.9 uM GGCAGGACAAAGAAGGTGGA; reverse, rP53-R/1, 0.9 uM, GGGCAGTGCTATGGAAGGAG; probe, rP53-P/1, 0.25 uM, [FAM]-CGCCCTTCAGCTTCACCCCA-ZEN/IBQ). The primers/probes and extracted DNA were mixed in a combined ddPCR reaction and run in standard conditions as described below. Percent chimerism was measured as either *rat* or *mouse* copies/ul divided by the sum of *mouse* and *rat* copies/ul.

For human>mouse chimeras, we previously designed a ddPCR assay to measure a wide range of human chimerism in mammalian tissues. This consisted of three multiplexed sets of primers and probes: (1) mammalian DNA reference which measures the total copies/ul of mammalian genomes (forward, Zeb2-F/5, 0.36 uM, GGATGGGGAATGCAGCTCTT; reverse, Zeb2-R/5.1, 0.36 uM, AGTGCGGCAGAATACAGCA; probe, Zeb2-P/5, 0.1 uM, [HEX]-TGATGGGTTGTGAAGGCAGCTGCACCT-ZEN/IBQ); (2) human gene specific target to measure the copies/ul of human genomes (forward, chCasp7-F/3.2, 0.9 uM, GAAGAAGAAAAATGTCACCATGCG; reverse, chCasp7-R/3.1, 0.9 uM, TTTCAAAATTCATGTTGTACTGATATGTAG; probe, chCasp7-P/3, 0.25 uM, [HEX]-AGACCACCCGGGACCGAGTGCCT-ZEN/IBQ); and another human specific primer/probe set that detects a repeat element (∼5000 repeats per cell) in the human genome to increase sensitivity in low level human>mouse chimeras (forward, AluYB8-F/2.1, 0.33 uM, GAAGAAGAAAAATGTCACCATGCG; reverse, AluYB8-R/2, 0.33 uM, TTTCAAAATTCATGTTGTACTGATATGTAG; probe, AluYB8-P/2.1, 0.09 uM, [FAM]-AGACCACCCGGGACCGAGTGCCT-ZEN/IBQ). The primers/probes and extracted DNA were mixed in a combined ddPCR reaction and run in standard conditions as described below. While AluYB8 was on the FAM channel, Zeb2 and chCasp7 were amplitude-multiplexed on the HEX channel such that Zeb2 was low and chCasp7 was high. For samples with a higher contribution of human cells, percent chimerism was measured as chCasp7 divided by Zeb2. For samples with a lower contribution of human cells, percent chimerism was measured as AluYB8 divided by 2500, divided by Zeb2 (note chCasp7 and Zeb2 have two copies per cell).

Each ddPCR reaction was prepared and analyzed with the QX200 ddPCR system (BioRad, Hercules, CA) in accordance with BioRad’s standard recommendations for use with their ddPCR Supermix for Probes (No dUTP). All reactions had a final volume of 20 ul loaded into the droplet generator, each containing 0.1-4 ul of crude lysate. Thermocycler conditions: 95 °C × 10 minutes; 50 cycles of 94 °C x 30 sec and 60 °C x 60 sec; 98 °C × 10 minutes; hold at 4°C. All primers and probes were ordered from Integrated DNA Technologies.

### Flow cytometry analysis of phagocytosis in intraspecies and interspecies chimeras

Single-cell suspensions were prepared from embryonic tissues of chimeras at E11.5 or E13.5. Cells were incubated with the indicated antibodies for 30 minutes at 4□°C in the dark. Dead cells were excluded using propidium iodide (PI, BioLegend, 421301). The following chimeric combinations were analyzed: E11.5 rat□>□mouse, E11.5 mouse□>□mouse, E13.5 rat□>□rat, E13.5 mouse□>□rat, and E11.5 human□>□mouse. For mouse host chimeras, macrophages were identified as CD45.1□CD11b□F4/80□ cells using the following antibodies:

- Mouse CD45.1 (PE, clone A20, BioLegend, 110707)
- Mouse CD11b (BUV395, clone M1/70, BD Biosciences, 563553)
- Mouse F4/80 (Pacific Blue, clone BM8, BioLegend, 123124)

For rat host chimeras, macrophages were defined as rat CD45□CD11b/c□ cells using:

- Rat CD45 (APC, clone OX-1, BioLegend, 202206)
- Rat CD11b/c (PE/Cy7, clone OX-42, BioLegend, 201806)

To quantify phagocytic activity, a “phagocytic index” was defined as the percentage of host macrophages (as above) containing detectable donor-derived fluorescent signals (EGFP or tdTomato), normalized to the total donor chimerism within each embryo. Index calculation was performed independently for each sample.

### Flow cytometry analysis of PtdSer by Annexin-V staining in intraspecies and interspecies chimeras

Single-cell suspensions were prepared from embryonic tissues of chimeras at E11.5 or E13.5 and stained to assess phosphatidylserine (PtdSer) exposure. The following chimeric combinations were analyzed: E11.5 rat□>□mouse, E11.5 mouse□>□mouse, E13.5 rat□>□rat, E13.5 mouse□>□rat, and E11.5 human□>□mouse.

Cells were washed twice with cold Annexin V Binding Buffer (BioLegend, 422201) and resuspended at a concentration of 1 × 10□ cells/mL, and 100□µL of cell suspension was transferred to each tube. Alexa Fluor® 647–conjugated Annexin V (BioLegend, 640911) was added at 5□µL per tube and incubated for 15 minutes at room temperature (25□°C) in the dark. After incubation, 400□µL of Annexin V Binding Buffer was added to each tube. Cells were stained with propidium iodide (PI, BioLegend, 421301) to exclude dead cells and analyzed immediately by flow cytometry.

### Comprehensive immune cell type profiling in E11.5 rat >mouse chimeras and adult mouse spleen

Single-cell suspensions were prepared from E11.5 rat□>□mouse chimeras and from spleens of 8-week-old female CD1 mice. Spleen samples were mechanically dissociated by gently sliding between glass slides, followed by red blood cell lysis using ACK lysis buffer (Gibco, A1049201). All samples were stained for 30 minutes at 4□°C in the dark using antibody cocktails diluted in staining buffer. Dead cells were excluded using propidium iodide (PI, BioLegend, 421301). To prevent fluorochrome interactions among Brilliant Violet dyes, Brilliant Stain Buffer (BD Biosciences, 563794) was used during antibody incubation. A list of monoclonal antibodies used for flow cytometry can be found in Table S3.

The following antibodies were used for immune cell subset profiling:

- Mouse CD11b (BUV395, clone M1/70, BD, 563553)
- Mouse F4/80 (BV421, clone BM8, BioLegend, 123124)
- Mouse CD45.1 (PE, clone A20, BioLegend, 110707)
- Mouse Ly6G (APC/Cy7, clone 1A8, BioLegend, 127624)
- Mouse Ly6C (PE/Cy7, clone HK1.4, BioLegend, 128017)
- Mouse B220/CD45R (Alexa Fluor 488, clone RA3-6B2, BioLegend, 103223)
- Mouse CD3 (BV605, clone 17A2, BioLegend, 100237)
- Mouse NK1.1 (APC, clone PK136, BioLegend, 108709)

Flow cytometry was performed immediately after staining using a BD FACSymphony™ analyzer.

### Flow cytometry analysis of macrophages in organs from PU.1 knockout or Csf1r knockout mice

Single-cell suspensions were prepared from brain, heart, lung, and spleen of E18.5 embryos derived from PU.1 heterozygous or Csf1r heterozygous intercrosses. Tissues were dissociated and passed through a 70□μm cell strainer to obtain single-cell suspensions. Cells were stained with fluorescently conjugated antibodies for 30 minutes at 4□°C in the dark. Dead cells were excluded using propidium iodide (PI, BioLegend, 421301).

The following antibodies were used to identify macrophage and myeloid populations:

- Mouse CD45.1 (PE, clone A20, BioLegend, 110707)
- Mouse CD11b (BUV395, clone M1/70, BD, 563553)
- Mouse Ly6G (APC/Cy7, clone 1A8, BioLegend, 127624)
- Mouse Ly6C (PE/Cy7, clone HK1.4, BioLegend, 128017)

### *In vitro* co-culture of mouse macrophages with heterogeneous cells from mouse or rat embryonic bodies

Mouse bone marrow-derived macrophages (BMDMs) were generated by culturing bone marrow cells from adult mice in Iscove’s Modified Dulbecco’s Medium (IMDM; Gibco, 12440053) supplemented with 10% fetal bovine serum (FBS), 1× MEM non-essential amino acids (NEAA; Gibco, 11140050), 1×penicillin-streptomycin (Gibco, 15140122), and 100□ng/mL recombinant mouse macrophage colony-stimulating factor (M-CSF; PeproTech, 315-02). Cultures were maintained for 7 days at 37□°C in 5% CO□. On day 7, BMDMs were primed with 100□ng/mL lipopolysaccharide (LPS from Escherichia coli O111:B4; Sigma-Aldrich, L4391) for 24 hours.

To mimic early-stage embryonic tissue, embryoid bodies (EBs) were generated from EGFP-expressing mouse or rat pluripotent stem cells using standard suspension culture protocols. On day 10, EBs were dissociated into single-cell suspensions and resuspended in serum-free IMDM.

For co-culture experiments, 1□×□10□ LPS-primed BMDMs were seeded in ultra-low attachment 96-well plates (Corning, 7007) and incubated with EB-derived single cells at an effector-to-target (E:T) ratio of 1:2 for 6 hours at 37□°C. Where indicated, BMS777607, a pan-inhibitor of the TAM receptor tyrosine kinase family (MedChemExpress, HY-10162), was added at a final concentration of 3□μM at the beginning of the co-culture.

After co-culture, cells were harvested and stained with antibodies against mouse CD11b (BUV395, clone M1/70, BD, 563553) and mosue F4/80 (BV421, clone BM8, BioLegend, 123124) to identify macrophages. Phagocytic activity was quantified by flow cytometry as the percentage of CD11b□F4/80□ macrophages containing EGFP□ donor-derived signals.

### Imaging cytometry analysis of “eating macrophages” from E11.5 rat > mouse chimeras

Single-cell suspensions were prepared from E11.5 rat□>□mouse chimeras. The rat cells were labeled with EGFP. Cells were incubated with the PE conjugated anti mouse CD45.1 antibody (BioLegend, 110707), APC conjugated anti mouse CD11b antibody (BioLegend, 101211), and Pacific Blue conjugated anti mouse F4/80 antibody (BioLegend, 123123) for 30 minutes at 4□°C in the dark. Dead cells were excluded using propidium iodide. Cells were resuspended in FACS buffer and analyzed by the “High-speed fluorescence image–enabled cell sorter (ICS)” using Cell View Technology^36^. “Eating macrophages” (mCD45.1^+^, CD11b^+^, F4/80^+^, EGFP^+^) and “Alive rat cells” (mCD45.1^-^, EGFP^+^) were selected for acquiring images. Bright field, EGFP and CD45.1 (PE channel) were imaged.

### Western blot analysis of Bcl2 expression level in six different cell lines

Cells were detached and lysed using RIPA buffer (1% Triton X-100, 1% Sodium Deoxycholate, 0.1% SDS, 150 nM NaCl, 10 mM Tris pH 8, 1x complete EDTA-free protease inhibitor cocktail (PIC), 1mM PMSF) for 10 minutes on ice. Lysates were then centrifuged at 16,000 x g for 10 minutes at 4°C. The supernatant was collected and quantified using BCA (Thermo Scientific, 23227). Following normalization, the protein was denatured by the addition of 1x NuPAGE LDS Sample Buffer (Invitrogen, NP0007) and 5% beta-mercaptoethanol (Sigma-Aldrich, M6250) and heating to 70°C for 10 minutes. 20 ug of protein per sample was then loaded onto a pre-cast 4-20% Novex Tris-glycine gel (Invitrogen, XP04200BOX) and run at 100V for 2.5 hours in Tris-glycine running buffer (25 mM Tris, 192 mM glycine, 0.1% SDS). The Gel was then transferred onto a nitrocellulose membrane (cytiva) for 1 hour at 400 mA in Tris-glycine transfer buffer (25 mM Tris, 192 mM glycine, 20% methanol). The membrane was stained with 0.1% Ponceau S in 3% trichloroacetic acid to assess overall transfer and relative protein levels between lanes. Then, the membrane was blocked in blocking solution (5% Blotto non-fat dry milk and 1% BSA in PBS with 0.1% Tween-20) for 30 minutes at room temperature on an orbital shaker, and then incubated with primary antibody in blocking solution overnight at 4°C. The blot was then washed 3x with PBS with 0.1% Tween-20 and incubated with horseradish peroxidase (HRP)-conjugated secondary antibody in blocking solution for 1 hour at room temperature. After 3x washes with PBS with 0.1% Tween-20, Amersham enhanced chemiluminescence Prime reagent (Cytiva, RPN2232) was used to perform chemiluminescence and imaged with an Amersham ImageQuant 800 (Amersham). The antibodies used include Bcl-2 (F9V5R) Rabbit mAb (Cell Signaling Technology, 28150, 1:1000), Oct-4A (C30A3) Rabbit mAb (Cell Signaling Technology, 2840, 1:1000), vinculin-HRP (Santa Cruz Biotechnology, sc-73614, 1:2000), Donkey anti-Rabbit IgG (H+L) HRP (Jackson Immunoresearch, 711-035-152, 1:3000).

### Single cell RNA sequencing of index sorted macrophages with Smart-seq

#### Single cell capture and RNA extraction

scRNA-seq with index sorting was performed as previously described^50^. Briefly, 96-well plates were prepared with 2 μL lysis buffer per well (1 U/μL RNase inhibitor (Clontech, 2313B), 0.1% Triton (ThermoFisher Scientific, 85111), 2.5 mM dNTP (Thermo Fisher Scientific, 10297018), 2.5 μM oligo dT_30_VN (Integrated DNA Technologies, 5′-AAGCAGTGGTATCAACGCAGAGTACT_30_VN-3’), and 1:600,000 ERCC (external RNA controls consortium) RNA spike-in mix at 1:600,000 (Thermo Fisher Scientific, 4456739) in UltraPure water (ThermoFisher Scientific, 10977015))^87^, which were stored at −80°C until use. Embryos were dissociated into a single cell suspension and stained for FACS as described above. Cells were sorted directly into lysis buffer plates on “single cell” precision mode with index sorting turned on. After a cell was sorted into each well, plates were centrifuged at 3000×*g* for 30 seconds at 4°C, immediately snap frozen on dry ice, then stored at −80°C.

#### Reverse transcription and pre-amplification

Reverse transcription (RT) and cDNA pre-amplification were carried out with a modified Smart-seq3 protocol^49^. First, lysis buffer plates were first incubated at 72°C for 3 minutes, then immediately snap chilled on ice in order to anneal the oligo dT_30_VN primer.

For RT, 3 μL of RT mix was added to each well (25 mM Tris-HCl pH 8.5 (Teknova, T5085), 0.5 U/μL RNase inhibitor (Clontech, 2313B), 8 mM dithiothreitol (DTT) (Promega, P1171), 30 mM NaCl (Thermo Fisher Scientific, AM9760G), 2.5 mM MgCl_2_ (Thermo Fisher Scientific, AM9530G), 1 mM GTP (ThermoFisher Scientific, R0461), 2 μM TSO (Integrated DNA Technologies, 5′-AAGCAGTGGTATCAACGCAGAGTGAATrGrGrG-3′), 5% polyethylene glycol (Sigma, P1458), and 2 U/μL Maxima H-minus reverse transcriptase (ThermoFisher Scientific, EP0753) in UltraPure water). Plates were thoroughly agitated to ensure mixture, spun down, then incubated in a C1000 Touch Thermal Cycler (BioRad) with the following program: (1) 42°C for 90 min for RT reaction, then (2) 70°C for 15 min to terminate the reaction.

For preamplification, 7.5 uL of PCR mix was added to each well (1.67X KAPA HiFi HotStart ReadyMix (Kapa Biosystems, KK2602) and 0.17 μM IS PCR primer (Integrated DNA Technologies, 5′-AAGCAGTGGTATCAACGCAGAGT-3′) in UltraPure water). Plates were again thoroughly agitated, spun down, then incubated using the following program: (1) 98°C for 3 min, (2) 98°C for 20 sec for denaturing, (3) 67°C for 15 sec for annealing, (4) 72°C for 6 minutes for elongation, (5) repeat from step 2 24 times, and (6) 72°C for 5 minutes.

Preamplified cDNA was purified to remove small oligos and residual reaction components. cDNA purification was performed by adding 0.65X volume of calibrated AMPure XP beads (Beckman Coulter, A63882) to each well, followed by two washes with 80% EtOH, and finally elution in 12.5 μL UltraPure water.

#### Quality control

Quality control was performed on the purified cDNA of each single cell on Fragment Analyzer (Advanced Analytical), a capillary electrophoresis-based machine. From the purified cDNA, 1 μL was taken from each single cell for analysis of concentration and size distribution. We set a minimum cDNA concentration cutoff of 1.7 ng/μL based on the cDNA concentration of empty wells containing ERCC but no sorted cell. Cells meeting the cDNA concentration cutoff were consolidated from 96 well plates into 384 well plates using a Mosquito X1 liquid handler (SPT Labtech). During consolidation, the cDNA concentration of each single cell was normalized to be within the 1.7–4.0 ng/μL range by diluting with an appropriate amount of UltraPure water.

#### Library preparation and sequencing

Library preparation steps were carried out in 384-well plate format using a Mosquito HTS liquid handler (SPT Labtech). Tagmentation was performed by combining 0.4 μL cDNA from each single cell with 1.2 μL Tn5 mix (1 ng/μL homebrewed Tn5 enzyme, 16 mM Tris-HCl pH 7.6, 16 mM MgCl_2_ (Thermo Fisher Scientific, AM9530G) and 8% dimethylformamide (DMF) (Thermo Fisher Scientific, AC327171000) in UltraPure water. The tagmentation reaction was incubated at 55°C for 10 min. The reaction was terminated by adding 0.4 μL neutralization buffer (0.1% SDS). Indexing PCR reactions were performed using a custom made 7680-plex unique dual-index primer set (Integrated DNA Technologies). 0.4 μL of 5 μM i5 indexing primer, 0.4 μL of 5 μM i7 indexing primer, and 1.2 μL KAPA HiFi HotStart ReadyMix (Kapa Biosystems, KK2602) was added to each well. Indexing PCR amplification was performed with the following program: (1) 72°C for 3 min, (2) 95°C for 30 sec, (3) 98°C for 10 sec for denaturing, (4) 67°C for 30 sec for annealing, (5) 72°C for 60 sec for elongation, (6) repeat from step 3 10 times. Following this amplification, 1 μL reaction product was taken from each well and pooled, forming a 384-cell pool. Each 384-cell library pool was purified using 0.8X volume of AMPure XP beads (Beckman Coulter, A63882), then analyzed on Fragment Analyzer (Advanced Analytical) for concentration and size distribution. Up to 20 384-cell library pools were normalized for concentration, pooled into a 7680-cell library pool, and purified again using 0.8X AMPure beads. The final 7680-cell library pool was sequenced on a NovaSeq 6000 S4 flow cell (Illumina) for 1-2 million 2×150 bp paired-end reads per cell.

#### Read mapping

Demultiplexing of reads was performed using bcl2fastq version 2.19.0.316. 3’ adapter sequences were removed using skewer v0.2.2^88^. Reads were then aligned to the mouse genome (GRCm39, mm39) as well as the rat genome (mRatBN7.2, rn7) using STAR version 2.6.1d using 2-pass mapping^89^. In the first pass, reads for each cell were aligned using a STAR genome index generated using the public transcript annotation for the mouse or rat genome (for mouse, Gencode version 29; for rat, release 110). After first pass mapping, mapped splice junctions from each cell were extracted and concatenated. A new STAR genome index was generated by combining the existing Gencode annotation with the newly discovered splice junctions from the first pass. Second pass mapping was then performed with this new STAR genome index which contains both known and newly identified splice junction. ENCODE recommended parameters (https://github.com/ENCODE-DCC/long-read-rna-pipeline) were used in addition to the option “--quantMode TranscriptomeSAM” during second-pass mapping to generate bam files for each cell. Gene and individual transcript expression levels were quantified from these bam files using RSEM version 1.3.3 with the “--single-cell-prior” setting^90^.

#### Data preprocessing

Gene count tables were concatenated from RSEM output, and integrated with metadata using Scanpy v.1.8.2^91^. Cells with <500 detected genes or <5000 read counts were filtered out. Genes expressed in fewer than 3 cells were filtered out. Read counts for each cell were normalized to 10,000, log transformed, and scaled to a maximum value of 10. Incidentally-sorted rat cells were identified and filtered out by comparing the percentage of uniquely mapped reads to the rat versus mouse genome. Highly-variable genes were identified using default parameters, which were then used to perform principle component analysis, compute the neighborhood graph, and cluster the cells with the Leiden method^92^. UMAP embeddings were computed to visualize data^93^. A total of 3065 cells across 12 embryos were included in the final analysis, with a median gene count of 7567 and read count of 5.97×10^5^. Step-by-step instructions to reproduce preprocessing and analysis of data are available on GitHub.

### Quantification and statistical analysis

For chimerism and phagocytic index comparison, different statistical tests were performed based on the biological replicate numbers in each group. When n ≥ 5, statistical significance was assessed using unpaired two tailed t-test, as indicated in the figures. When n < 5, statistical significance was assessed using unpaired Mann-Whitney U non-parametric test, as indicated in the figures. For comparing Annexin-V signal between donor and host cells within the same chimeric embryo, paired two tailed t-tests were performed, as indicated in Figure 4B. p-values were calculated using Prism 10 software. Flow cytometry data were analyzed using FlowJo v.10.10.0.

## Data availability

Single cell RNA-seq data is available on Synapse (SynID: syn68348899).

## Summary of embryo manipulation results in this study

A detailed summary of embryo manipulation results including embryo transfer numbers, implantation site numbers, embryo recovery rate and chimera frequency are provided in Table S4.

---

**Table S1.**
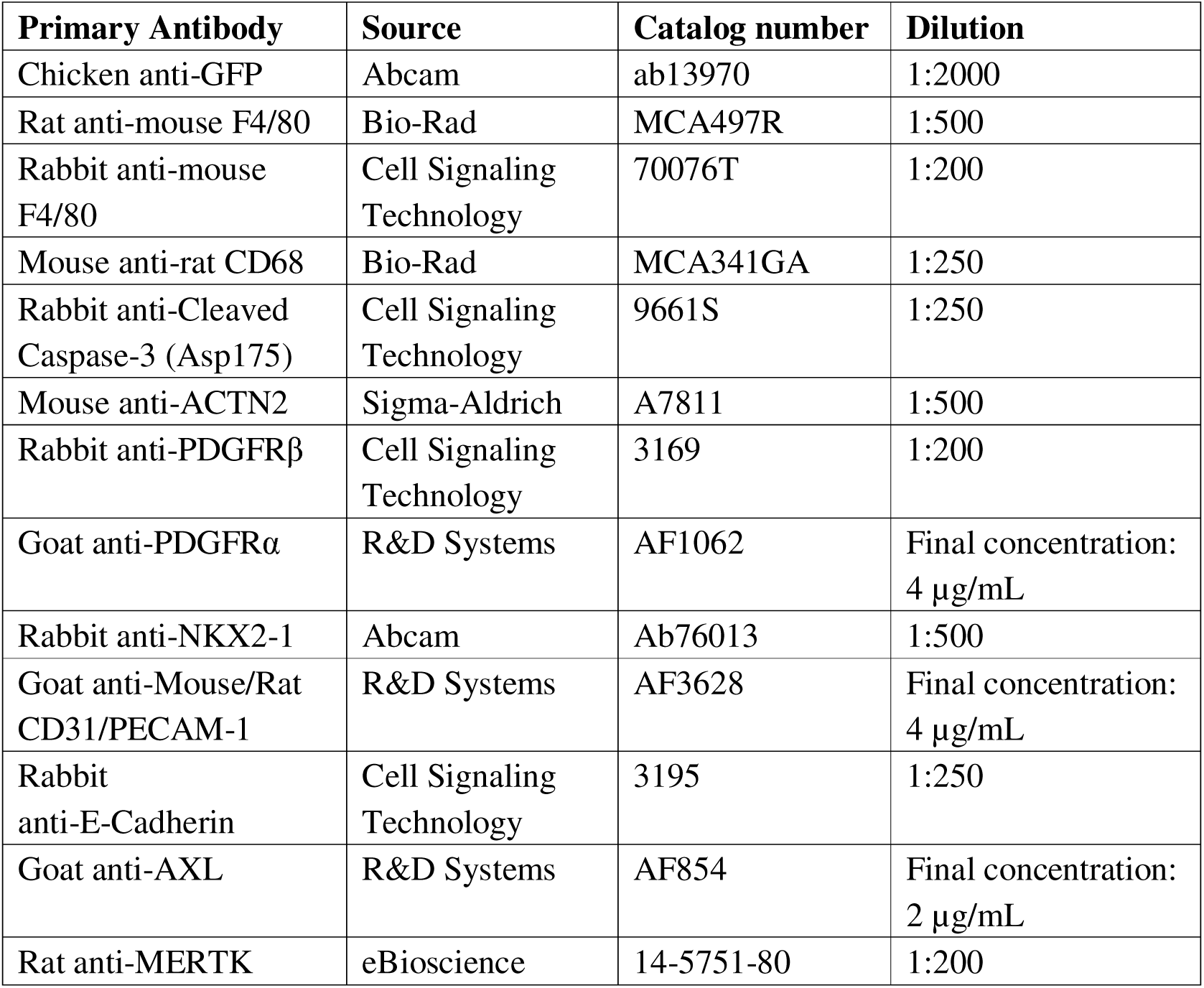
Primary antibodies used for immunohistochemistry in this study and their sources and dilutions.

**Table S2.**
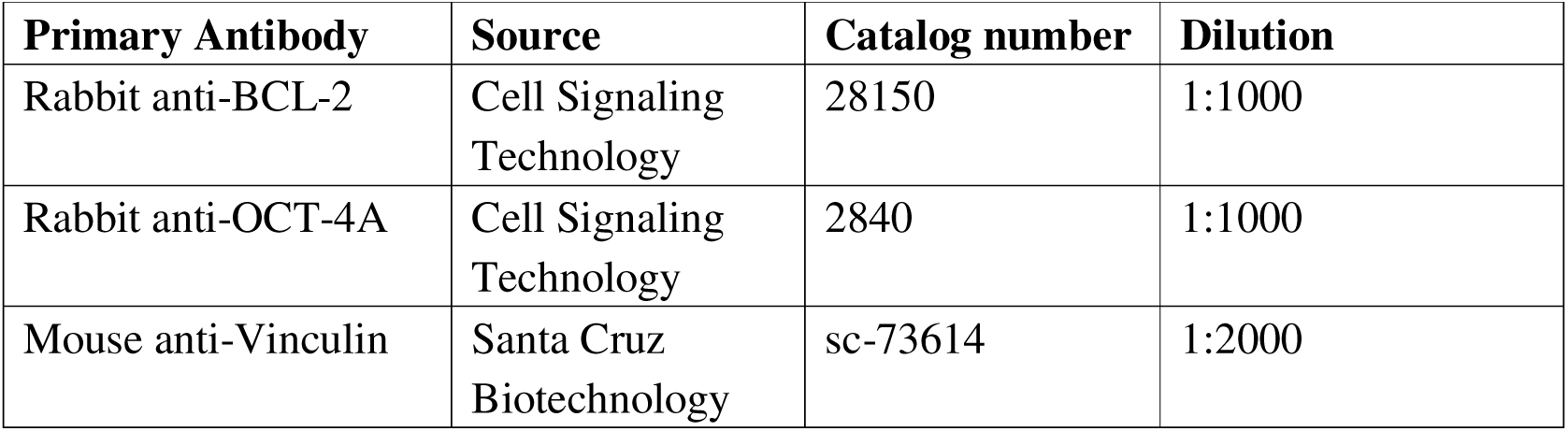
Primary antibodies used for western blot in this study and their sources and dilutions.

**Table S3.** Monoclonal antibodies used for flow cytometry analysis in this study and their sources and clone name.

| Antibody | Source | Catalog number | clone |
| --- | --- | --- | --- |
| Mouse CD45.1-PE | BioLegend | 110707 | A20 |
| Mouse CD11b-BUV395 | BD Biosciences | 563553 | M1/70 |
| Mouse CD11b-APC | BioLegend | 101211 | M1/70 |
| Mouse F4/80-Pacific Blue | BioLegend | 123124 | BM8 |
| Rat CD45-APC | BioLegend | 202206 | OX-1 |
| Rat CD11b/c-PE/Cy7 | BioLegend | 201806 | OX-42 |
| Mouse Ly6G-APC/Cy7 | BioLegend | 128017 | 1A8 |
| Mouse Ly6C-PE/Cy7 | BioLegend | 128017 | HK1.4 |
| Mouse B220-Alexa Fluor 488 | BioLegend | 103223 | RA3-6B2 |
| Mouse CD3-BV605 | BioLegend | 100237 | 17A2 |
| Mouse NK1.1-APC | BioLegend | 108709 | PK136 |

**Table S4.** Summary of embryo manipulation results in this study.

**Related to Figure 1A**
| Donor cell | Number of injected cells | Host embryo strain | Stage of analysis | Transferred embryos | Implantation <sup>(a)</sup> | Live embryos <sup>(b)</sup><br>(b/a%) | Chimeras <sup>(c)</sup><br>(c/b%) |
| --- | --- | --- | --- | --- | --- | --- | --- |
| Rat iPSC (T1-3) | 4-6 | CD1 | E9.5 | 60 | 42 | 26 (62%) | 14 (54%) |
| Rat iPSC (T1-3) | 4-6 | CD1 | E11.5 | 120 | 94 | 48 (51%) | 20 (42%) |
| Rat iPSC (T1-3) | 4-6 | CD1 | E14.5 | 60 | 41 | 17 (41%) | 6 (35%) |

**Related to Figure 2B**
| Donor cell | Number of injected cells | Host embryo strain | Stage of analysis | Transferred embryos | Implantation <sup>(a)</sup> | Live embryos <sup>(b)</sup><br>(b/a%) | Chimeras <sup>(c)</sup><br>(c/b%) |
| --- | --- | --- | --- | --- | --- | --- | --- |
| Rat iPSC (T1-3) | 4-6 | Wild-type CD1 | E9.5 | 30 | 20 | 15 (75%) | 9 (60%) |
| Rat iPSC (T1-3) | 4-6 | PU.1 KO CD1 | E9.5 | 60 | 40 | 20 (50%) | 14 (70%) |
| Rat iPSC (T1-3) | 4-6 | Wild-type CD1 | E11.5 | 100 | 77 | 34 (44%) | 20 (59%) |
| Rat iPSC (T1-3) | 4-6 | PU.1 KO CD1 | E11.5 | 110 | 60 | 25 (42%) | 16 (64%) |
| Rat iPSC (T1-3) | 4-6 | Wild-type CD1 | E14.5 | 48 | 37 | 16 (43%) | 6 (38%) |
| Rat iPSC (T1-3) | 4-6 | PU.1 KO CD1 | E14.5 | 60 | 39 | 14 (36%) | 7 (50%) |

**Related to Figure 2D**
| Donor cell | Number of injected cells | Host embryo strain | Stage of analysis | Transferred embryos | Implantation <sup>(a)</sup> | Live embryos <sup>(b)</sup><br>(b/a%) | Chimeras <sup>(c)</sup><br>(c/b%) |
| --- | --- | --- | --- | --- | --- | --- | --- |
| Rat iPSC (T1-3) | 4-6 | CD1 from PU.1 heterozygous intercrosses | E18.5 | 280 | - | 75 | 28 (37%) |

Related to Figure 2E
| Donor cell | Number of injected cells | Host embryo strain | Stage of analysis | Transferred embryos | Implantation <sup>(a)</sup> | Live embryos <sup>(b)</sup><br>(b/a%) | Chimeras <sup>(c)</sup><br>(c/b%) |
| --- | --- | --- | --- | --- | --- | --- | --- |
| Rat iPSC (T1-3) | 4-6 | CD1 from Csf1r heterozygous intercrosses | E18.5 | 180 | - | 46 | 19 (41%) |

Related to Figure 2F
| Donor cell | Number of injected cells | Host embryo strain | Stage of analysis | Transferred embryos | Implantation <sup>(a)</sup> | Live embryos <sup>(b)</sup><br>(b/a%) | Chimeras <sup>(c)</sup><br>(c/b%) |
| --- | --- | --- | --- | --- | --- | --- | --- |
| Mouse ESC (SGE2) | 6-8 | Wild-type CD1 | E18.5 | 25 | - | 12 | 10 (83%) |
| Mouse ESC (SGE2) | 6-8 | PU.1 KO CD1 | E18.5 | 15 | - | 6 | 5 (83%) |
| Mouse ESC (SGE2) | 6-8 | Csf1r KO CD1 | E18.5 | 15 | - | 6 | 6 (100%) |

Related to Figure 3G
| Donor cell | Number of injected cells | Host embryo strain | Stage of analysis | Transferred embryos | Implantation <sup>(a)</sup> | Live embryos <sup>(b)</sup><br>(b/a%) | Chimeras <sup>(c)</sup><br>(c/b%) |
| --- | --- | --- | --- | --- | --- | --- | --- |
| Rat iPSC (T1-3) | 6-7 | Wild-type CD1 | E11.5 | 100 | 58 | 30 (52%) | 17 (57%) |
| Rat iPSC (T1-3) | 6-7 | Axl KO CD1 | E11.5 | 67 | 44 | 17 (39%) | 9 (53%) |
| Rat iPSC (T1-3) | 6-7 | Mertk KO CD1 | E11.5 | 35 | 26 | 12 (46%) | 4 (33%) |
| Rat iPSC (T1-3) | 6-7 | Axl, Mertk double KO CD1 | E11.5 | 60 | 41 | 13 (32%) | 8 (62%) |

Related to Figure 4C
| Donor cell | Number of injected cells | Host embryo strain | Stage of analysis | Transferred embryos | Implantation <sup>(a)</sup> | Live embryos <sup>(b)</sup><br>(b/a%) | Chimeras <sup>(c)</sup><br>(c/b%) |
| --- | --- | --- | --- | --- | --- | --- | --- |
| Control rat iPSC with EGFP OE | 5-6 | Wild-type CD1 | E18.5 | 80 | - | 17 | 7 (41%) |
| Rat iPSC with BCL2 OE | 5-6 | Wild-type CD1 | E18.5 | 80 | - | 22 | 14 (64%) |
| Rat iPSC with<br>mouse CD47 OE | 5-6 | Wild-type CD1 | E18.5 | 80 | - | 19 | 10 (53%) |

**Related to Figure 5H**
| Donor cell | Number of<br>injected cells | Host embryo<br>strain | Stage of<br>analysis | Transferred<br>embryos | Implantation <sup>(a)</sup> | Live embryos <sup>(b)</sup><br>(b/a%) | Chimeras <sup>(c)</sup><br>(c/b%) |
| --- | --- | --- | --- | --- | --- | --- | --- |
| Human iPSC<br>(CAG-BCL2, TRE-MYC) | 4-5 | Wild-type CD1 | E11.5 | 60 | 41 | 24 (59%) | 11 (46%) |
| Human iPSC<br>(CAG-BCL2, TRE-MYC) | 4-5 | PU.1 KO CD1 | E11.5 | 40 | 24 | 10 (42%) | 6 (60%) |

**Related to Figure S5C**
| Donor cell | Number of<br>injected cells | Host embryo<br>strain | Stage of<br>analysis | Transferred<br>embryos | Implantation <sup>(a)</sup> | Live embryos <sup>(b)</sup><br>(b/a%) | Chimeras <sup>(c)</sup><br>(c/b%) |
| --- | --- | --- | --- | --- | --- | --- | --- |
| Human iPSC<br>(CAG-BCL2, TRE-MYC) | 4-5 | Wild-type CD1 | E9.5 | 30 | 24 | 18 (75%) | 7 (39%) |
| Human iPSC<br>(CAG-BCL2, TRE-MYC) | 4-5 | Wild-type CD1 | E11.5 | 37 | 32 | 25 (78%) | 7 (28%) |

## ACKNOWLEDGMENTS

We thank S. Homma for laboratory and administrative support; T. Nishimura for his generous training and advice on embryo work; T. Kobayashi for advice on rat ESC derivation and culture; S. Hamanaka for her advice on interspecies chimera assays; C. Carswell-Crumpton and C. Pan for their advice on flow cytometric assays; A. Barkal and A. Banuelos for their advice on macrophage phagocytosis assays; K. Alvarez, M.R. Eckart, and the Stanford Protein and Nucleic Acid (PAN) Facility for technical support with cDNA quality control; A.H. Chang, R. Fueyo, T. Chern, L. Ichino and M. Koska for their critical reading of the manuscript. Stanford Stem Cell Institute FACS Core provided access to flow cytometers. Stanford Transgenic, Knockout and Tumor model Center provided assistance for preparing pseudo pregnant mice.

This research was supported by the grants from the NIH (R01DK121851), the Ludwig Foundation, the Leducq Foundation, the Centers for Clinical Application Research on Specific Disease/Organ of the Research Center Network for Realization of Regenerative Medicine, funded by the Japan Agency for Medical Research and Development (AMED_JP23bm1123041 to H.N.) and the Japan Society of the Promotion of Science.

S.W. is supported by Stanford Graduate Fellowship and CIRM training grant. K.N. was supported by a research fellowship from Japan Society for the Promotion of Science (JSPS). D.D.L is supported by Stanford University Medical Scientist Training Program grant T32-GM007365 and T32-GM145402, and the Seth A. Ritch Bio-X Stanford Interdisciplinary Graduate Fellowship (SIGF). M.M. was supported by Uehara Memorial Foundation Overseas Research Fellowship and Stanford Cancer Institute Fellowship. I.L.W. is supported by an NIH/NCI Outstanding Investigator Award R35-CA220434 and the Virginia and D.K. Ludwig Fund for Cancer Research.

## AUTHOR CONTRIBUTIONS

S.W., K.N. and D.D.L conceived the research, performed the experiments, analyzed the data and wrote the manuscript. S.W. designed and performed embryo manipulation, chimera generation and chimerism quantification. K.N. designed and performed flow cytometry analysis, immunology related assays and analyzed data. D.D.L designed and performed single cell RNA-seq experiments and bioinformatics analysis. F.S. developed the ddPCR assays for quantifying intraspecies and interspecies chimerism. F.S. supervised chimerism analysis using ddPCR data. H.S., A.Y. and H.M. generated the engineered hiPSC line and validated their interspecies chimera formation competency. M.M., S.T., N.H., J.B., C.T.C. and J.Z. performed experiments and analyzed data. I.L.W. supervised the experiments and participated in the interpretation of data. H.N. conceived the research, supervised the experiments, analyzed the data, and wrote the manuscript. All authors edited the manuscript.

## Supplemental figure legend

**Figure S1.**
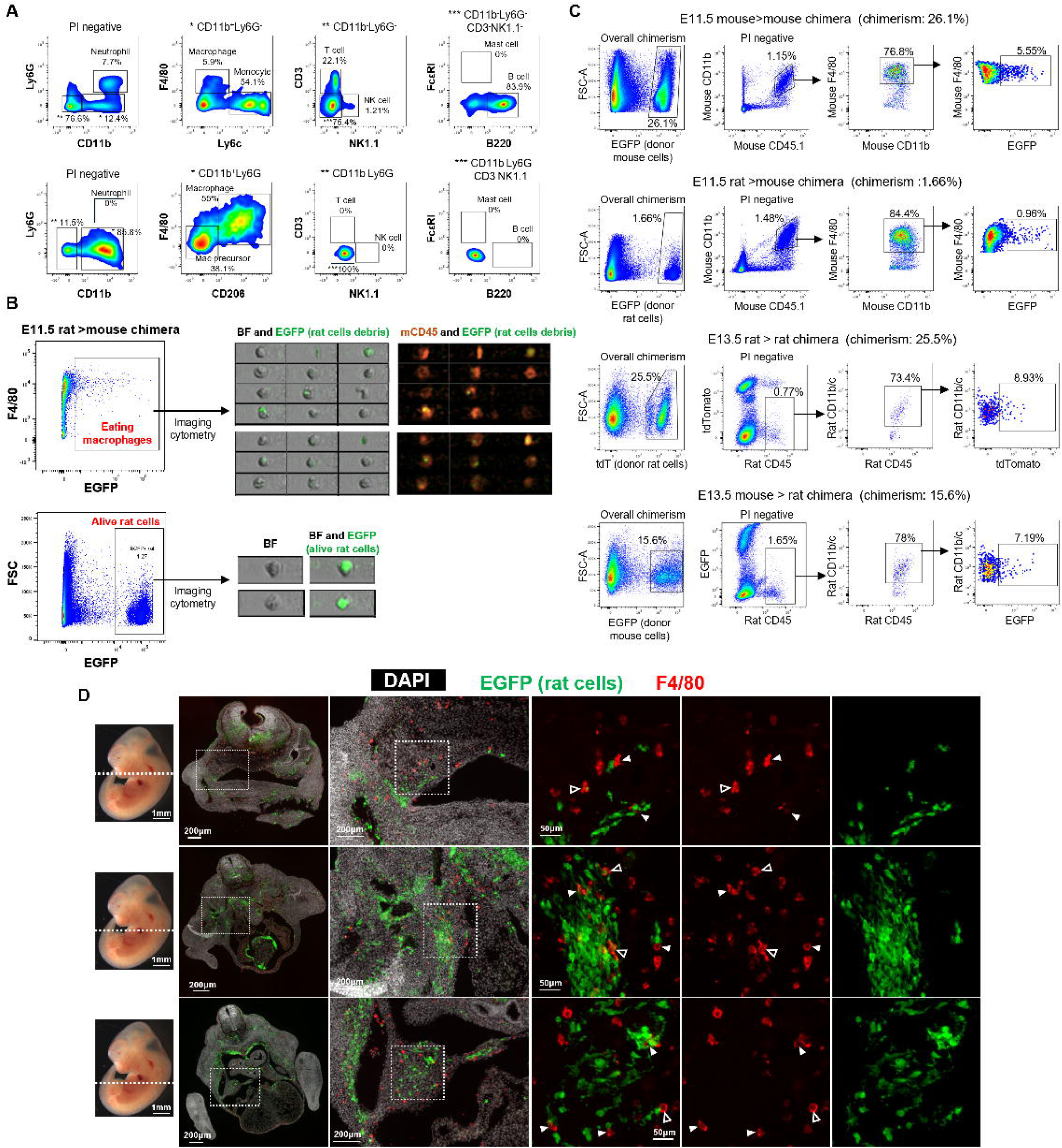
Quantification of phagocytosis in intraspecies and interspecies chimeras through flow cytometry and microscopy, related to Figure 1. **(A)** Flow cytometry gating strategy for detecting different types of CD45^+^ immune cells from adult CD1 splenocytes (top) or E11.5 rat > mouse chimera (bottom). The markers for differnt immune cells are: Neutrophil: CD11b^+^, Ly6G^+^; Adult macrophage: CD11b^+^, Ly6G^-^, F4/80^+^; Embryonic macrophage: CD11b^+^, Ly6G^-^, CD206^+^, F4/80^+^; Monocyte: CD11b^+^, Ly6G^-^, Ly6C^+^; Natural killer (NK) cells: CD11b^-^, Ly6G^-^, NK1.1^+^; CD3^+^ T cells: CD11b^-^, Ly6G^-^, CD3^+^; B cells: CD11b^-^, Ly6G^-^, CD3^-^, NK1.1^-^, B220^+^; Mast cells: CD11b^-^, Ly6G^-^, CD3^-^, NK1.1^-^, FcεRI^+^. **(B)** Representative imaging cytometry images showing “eating macrophages” (top) containing punctate EGFP signal and membrane-localized CD45, and an intact donor rat cell (bottom) with uniformly distributed cytosolic EGFP. **(C)** Flow cytometry gating strategy for detecting eating macrophages in four types of chimeras (top to bottom: E11.5 mouse> mouse, E11.5 rat> mouse, E13.5 rat>rat and E13.5 mouse>rat chimeras). The mouse macrophages are defined as CD45.1^+^, CD11b^+^, F4/80^+^ population. The rat macrophages are defined as rat CD45^+^, rat CD11b/c^+^ population. **(D)** Transverse sections of E11.5 rat□>□mouse chimeras stained for EGFP (rat donor cells) and F4/80 (mouse macrophages). Approximate sectioning planes are indicated by dashed lines on a schematic embryo. Stitched low-magnification images and corresponding high-magnification views are shown. Filled arrowheads mark macrophages in contact with donor cells (“contacting mode”); open arrowheads indicate macrophages containing internalized donor material (“phagocytizing mode”).

**Figure S2.**
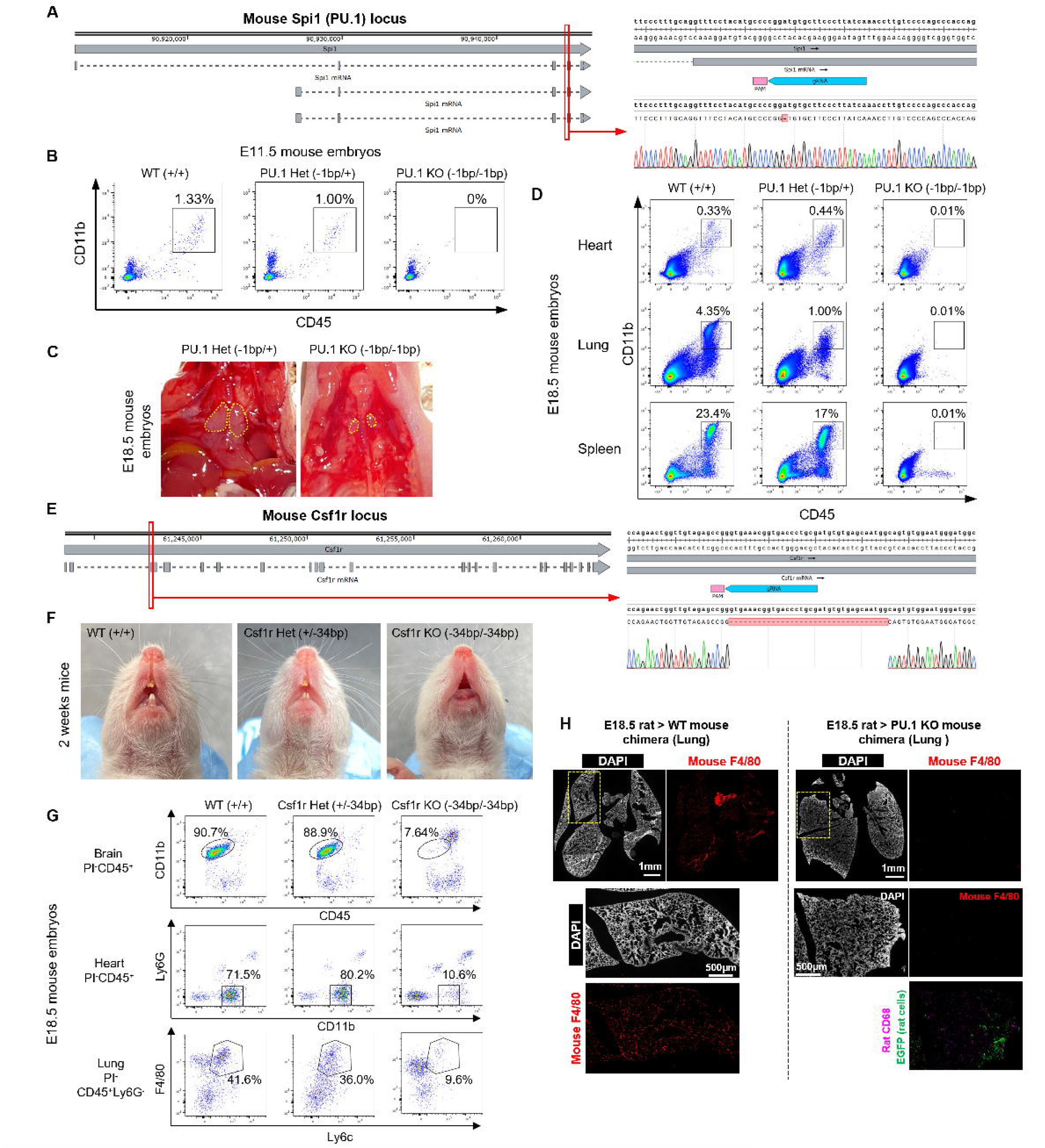
Phenotypic characterization of PU.1 and Csf1r knockout mice and chimeras, related to Figure 2. **(A)** Genotyping result of PU.1 knockout mice generated via CRISPR–Cas9 genome editing, showing a 1 bp deletion at the beginning of exon 5. **(B)** Flow cytometry plots of E11.5 embryos from PU.1 heterozygous intercrosses, showing WT, heterozygous, and knockout genotypes. Macrophages were identified using CD45 and Cd11b. **(C)** Gross morphology of thymus from E18.5 PU.1 heterozygous and knockout embryos. Yellow dashed lines outline the embryonic thymus region. **(D)** Flow cytometry analysis of immune cell composition in multiple organs from E18.5 embryos derived from PU.1 heterozygous intercrosses. **(E)** Genotyping result of Csf1r knockout mice generated via CRISPR–Cas9 genome editing, showing a 34 bp deletion at the beginning of exon 2. **(F)** Oral view of two-week-old WT, heterozygous, and Csf1r knockout mice. **(G)** Flow cytometry analysis of immune cell composition in multiple organs from E18.5 embryos derived from Csf1r heterozygous intercrosses. **(H)** Lung tissue sections from E18.5 rat□>□mouse chimeras with WT or PU.1 knockout hosts stained for EGFP (rat donor cells), F4/80 (mouse macrophages), and rat CD68 (rat macrophages).

**Figure S3.**
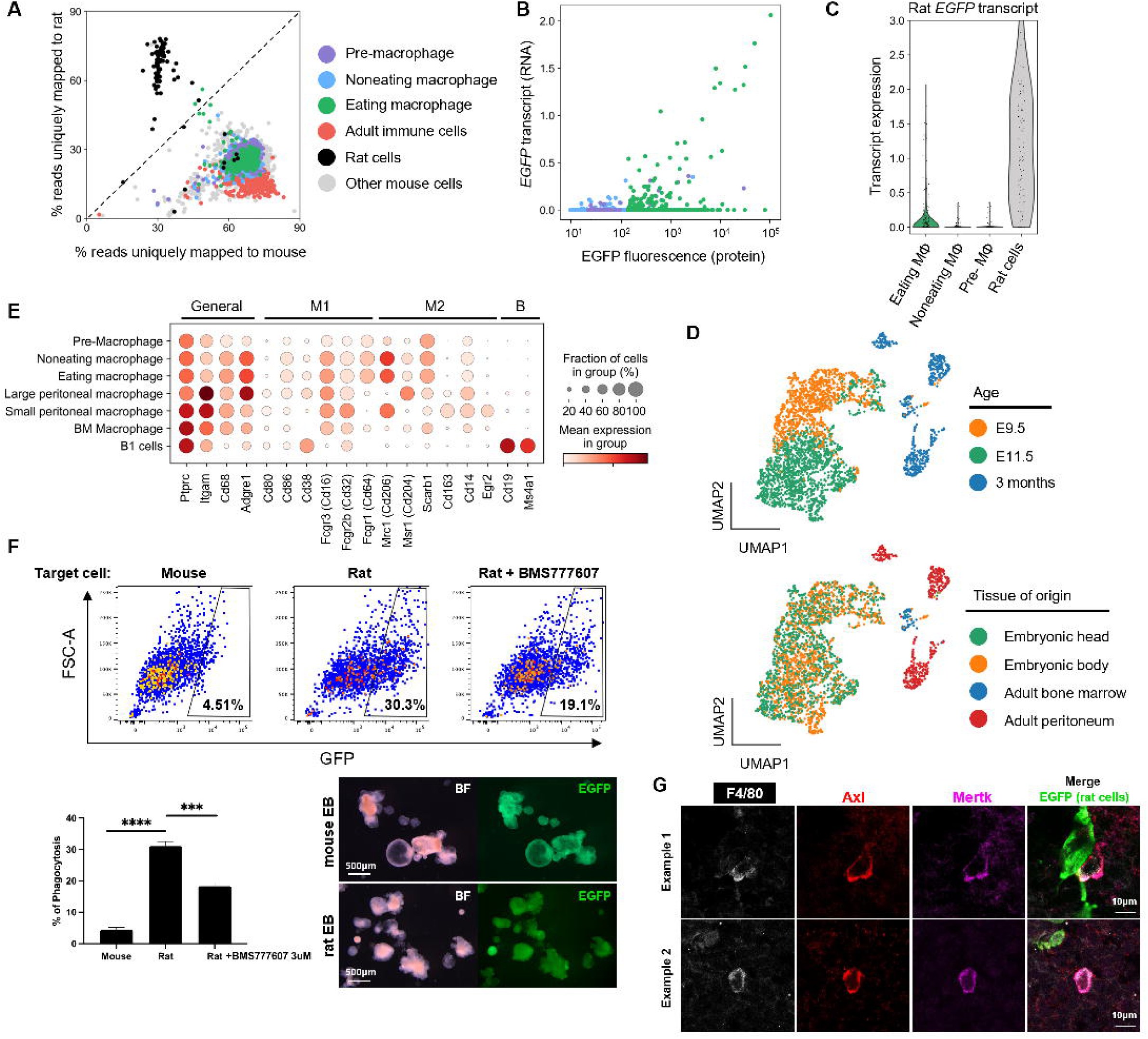
Single-cell characterization of macrophages from E9.5 and E11.5 rat□>□mouse chimeras, related to Figure 3. **(A)** Species assignment of index-sorted single cells based on the percentage of reads uniquely mapped to mouse or rat genomes. **(B)** Correlation of EGFP transcript with EGFP fluorescence intensity as measured by index sort from three macrophage populations shown in A. **(C)** EGFP transcript expression across four sorted populations: rat donor cells, eating macrophages, non-eating macrophages, and pre-macrophages. **(D)** UMAP of index-sorted cells colored by developmental stage and tissue of origin. **(E)** Dot plot showing the expression of selected M1 and M2 macrophage marker genes across sorted macrophage populations. **(F)** *In vitro* phagocytosis assay using mouse macrophages co-cultured with dissociated single cells from mouse or rat embryoid bodies, with or without the pan-TAM inhibitor BMS777607. Top: representative flow cytometry plots showing the percentage of macrophages undergoing phagocytosis. Bottom: quantification from two biological replicates. Statistical significance was assessed using unpaired two tailed t-test. ***p < 0.001, ****p < 0.0001. Representative images of EGFP-labeled mouse and rat embryoid bodies (day 10) are also shown. **(G)** Confocal images of F4/80□ macrophages in E11.5 rat□>□mouse chimeras stained for Axl and Mertk. Co-expression of Axl and Mertk is observed.

**Figure S4.**
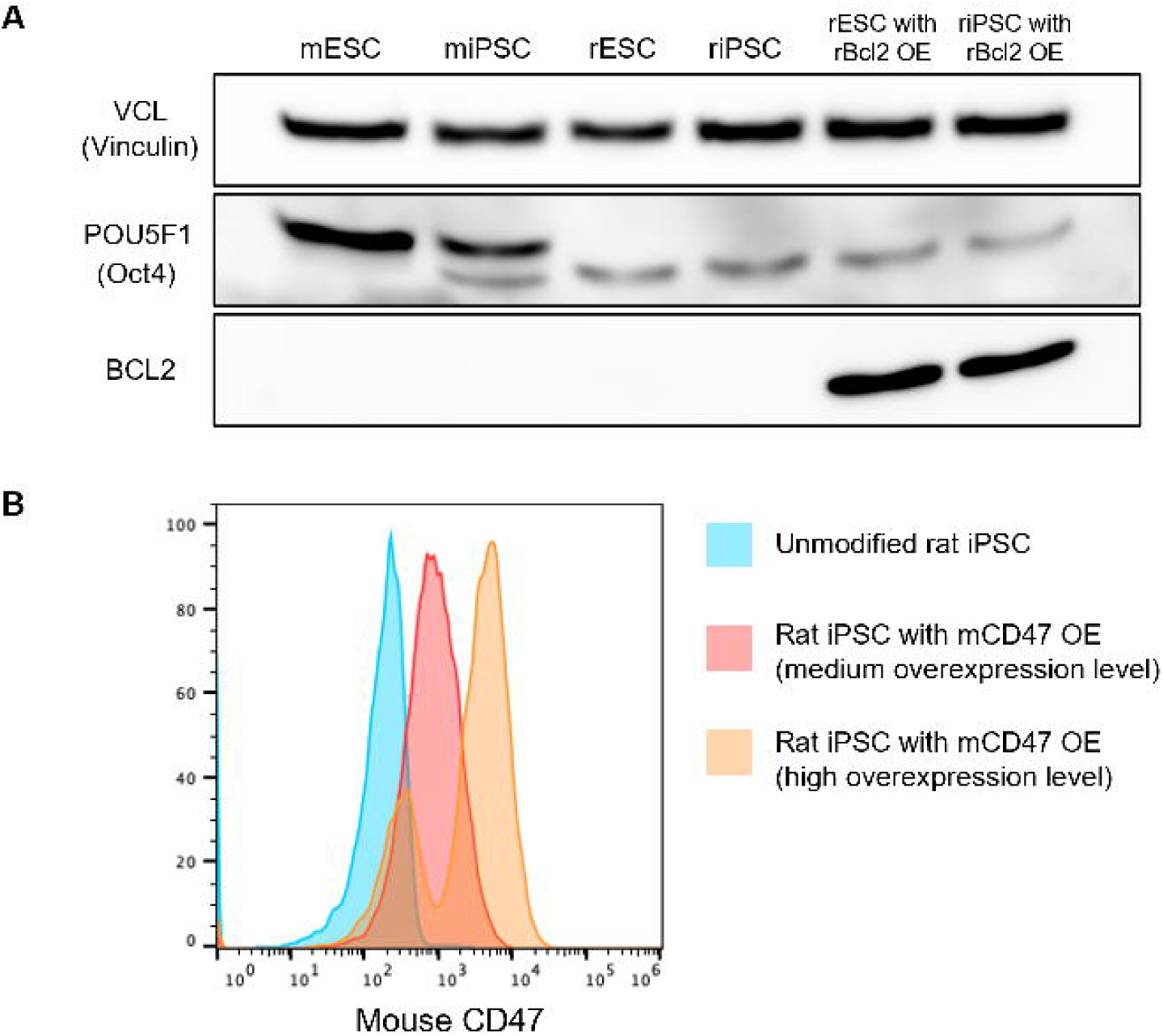
Generation and validation of rat pluripotent stem cell lines overexpressing rat Bcl2 or mouse CD47, related to Figure 4. **(A)** Western blot analysis of six cell lines using antibodies against Bcl2, Oct4 (pluripotency marker), and Vinculin (housekeeping control). Samples from left to right: C57B6 mouse ESCs, C57B6 mouse iPSCs, Wistar rat ESCs, Wistar rat iPSCs, rat Bcl2-overexpressing rat ESCs, and rat Bcl2-overexpressing rat iPSCs. **(B)** Flow cytometry histogram of mouse CD47 expression in unmodified and mCD47-overexpressing rat iPSCs.

**Figure S5.**
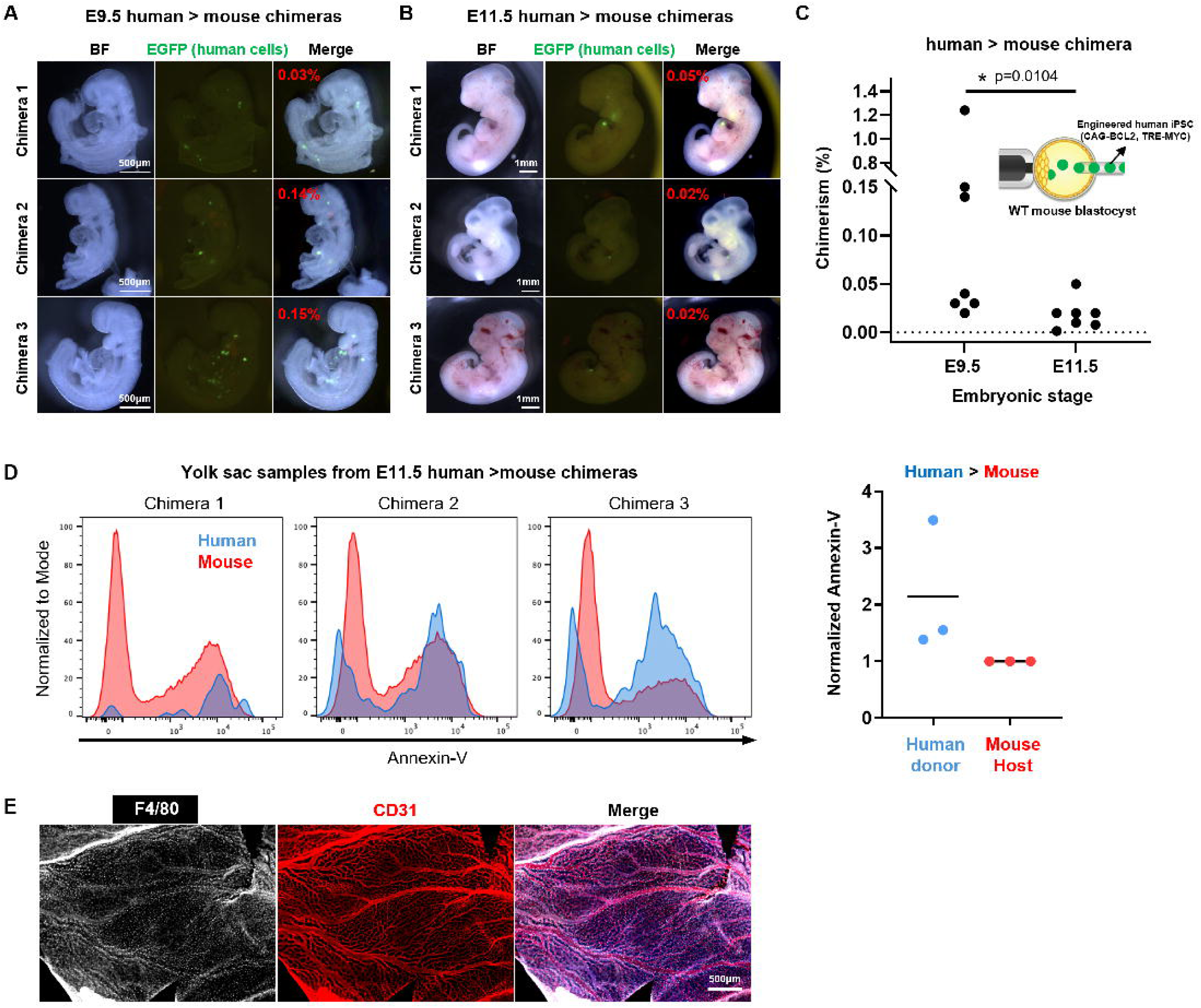
Characterization of human□>□mouse chimeras at E9.5 and E11.5, related to Figure 5. **(A-B)** Bright-field and fluorescence images of E9.5 (A) and E11.5 (B) human□>□mouse chimeras. Three chimeras per stage are shown. EGFP indicates human donor cells. ddPCR-determined chimerism levels are overlaid in red. **(C)** Whole-embryo chimerism analysis of human□>□mouse chimeras at E9.5 (n□=□7) and E11.5 (n□=□7). Each dot represents one chimeric embryo. Chimerism was quantified by ddPCR. Statistical significance was assessed using unpaired Mann-Whitney U non-parametric test. n.s., not statistically significant; *p < 0.05. **(D)** Left: flow cytometry histograms of Annexin-V staining in yolk sac samples from E11.5 human□>□mouse chimeras. Traces represent donor human cells (blue) and host mouse cells (red). Right: quantification of Annexin-V signal, shown as donor-to-host MFI ratios (n□=□3). **(E)** Fluorescence images of E11.5 mouse yolk sac stained for F4/80 (macrophages) and CD31 (endothelial cells).

**Figure S6.**
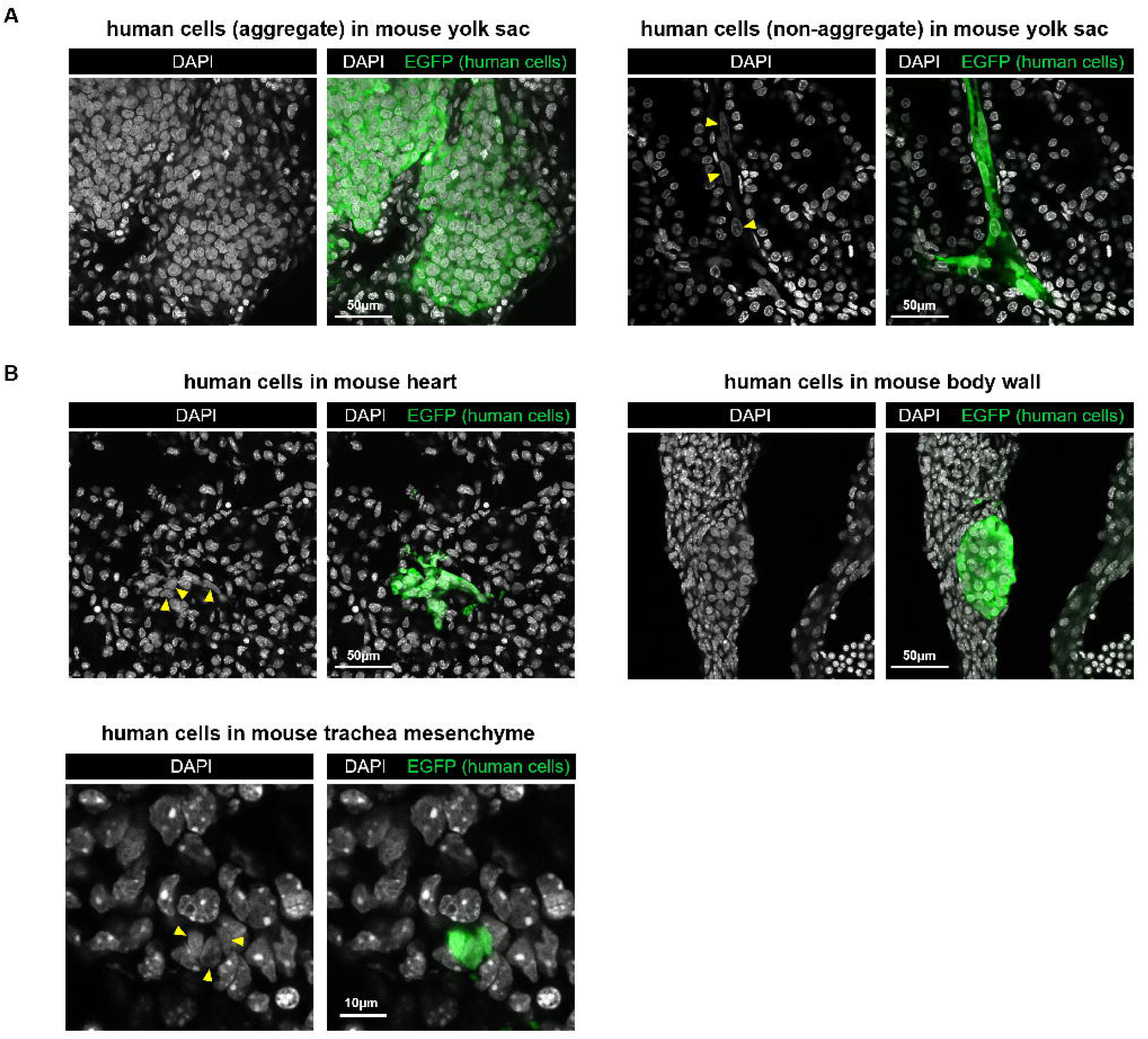
Human and mouse cells display different nuclear DNA staining patterns in human > mouse chimeras, related to Figure 5. **(A)** Confocal fluorescence images of human cells in the mouse yolk sac from E11.5 human > mouse chimeras. Yellow arrowheads highlight human nuclei with uniform DAPI signal. **(B)** Confocal fluorescence images of human cells identified in different regions of the mouse embryo from E11.5 human>mouse chimeras. Yellow arrowheads highlight human nuclei with uniform DAPI signal.

